# NLRP3 Cys126 palmitoylation by ZDHHC7 Promotes Inflammasome Activation

**DOI:** 10.1101/2023.11.07.566005

**Authors:** Tao Yu, Dan Hou, Jiaqi Zhao, Xuan Lu, Wendy K. Greentree, Qian Zhao, Min Yang, Don-Gerard Conde, Maurine E. Linder, Hening Lin

## Abstract

NACHT-, leucine-rich-repeat- (LRR), and pyrin domain-containing protein 3 (NLRP3) mediates inflammasome activation in response to multiple pathogen and damage-associated molecular patterns in macrophages. Hyperactivation of NLRP3 inflammasome contributes to many human chronic inflammatory diseases. Understanding how NLRP3 inflammasome is regulated can potentially provide new strategies to treat inflammatory diseases. Here, we demonstrated that NLRP3 is palmitoylated on Cys126 by palmitoyl-acyltransferase ZDHHC7 in macrophages, which is critical for NLRP3-mediated inflammasome activation. Perturbating NLRP3 Cys126 palmitoylation by ZDHHC7 knockout, pharmacological inhibition, or modification site mutation, diminishes NLRP3 activation and the consequential Caspase-1 and Gasdermin D (GSDMD) cleavage, and IL-1β and IL-18 secretion in mouse primary macrophages and human macrophages. Furthermore, NLRP3 Cys126 palmitoylation is vital for inflammasome activation *in vivo*, as *Zdhhc7* knockout, pharmacological inhibition, or NLRP3 C126A mutation protects mice from LPS-induced endotoxic shock and monosodium urate (MSU)-induced peritonitis. Mechanistically, ZDHHC7-mediated NLRP3 Cys126 palmitoylation promotes resting NLRP3 localizing on the *trans*-Golgi network (TGN) and activated NLRP3 on the dispersed TGN (dTGN), which is indispensable for the recruitment and oligomerization of adaptor protein ASC after inflammasome activation. The activation of NLRP3 by ZDHHC7-mediated Cys126 palmitoylation is different from the previously reported inhibitory effect by ZDHHC12-mediated Cys841 palmitoylation, highlighting the versatile regulatory roles of *S*-palmitoylation. Therefore, our study identifies a new regulatory mechanism of NLRP3 activation and suggests targeting ZDHHC7 or NLRP3 Cys126 residue as a potential therapeutic strategy to treat NLRP3-related human disorders.

**Highlights:**

- NLRP3 Cys126 is palmitoylated by ZDHHC7.
- ZDHHC7 promotes NLRP3 activation in macrophages, which can be inhibited by ZDHHCs inhibitors, 2-bromopalmitate and MY-D4.
- Cys126 palmitoylation of NLRP3 is critical for NLRP3 activation.
- NLRP3 TGN/dTGN localization depends on ZDHHC7-mediated Cys126 palmitoylation, which is crucial for ASC recruitment and inflammasome assembly.
- NLRP3 inflammasome activation by ZDHHC7 differs from ZDHHC12-mediated NLRP3 inhibition.

## Introduction

Macrophages play important roles in defending hosts from pathogen infection or sensing intracellular dangerous molecules. The pattern recognition receptors (PRRs) in macrophages recognize diverse pathogen-associated molecular patterns (PAMPs)^1^ or damage-associated molecular patterns (DAMPs)^2^ and promote pro-inflammatory responses by secreting cytokines such as interleukin-1β (IL-1β). NOD-like Receptor Protein 3 (NLRP3) is a cytosolic PRR that responds to multiple PAMPs/DAMPs and forms a multi-component complex called NLRP3 inflammasome to promote inflammation^3^. The activation of NLRP3 inflammasome involves a priming step and an activation step^4^. In the priming step, lipopolysaccharides (LPS, a component of Gram-negative bacterial cell wall) or other signals are recognized by cell surface PRRs and activate NF-κB signaling to induce expression of NLRP3, pro-IL-1β, and pro-IL-18^3,4^. During the activation step, multiple PAMPs or DAMPs, including the bacterial pore-forming toxin nigericin and extracellular adenosine triphosphate (ATP), induce cellular changes, such as potassium efflux and mitochondrial reactive oxygen species production, which activate NLRP3^3–5^. Activated NLRP3 recruits ASC (apoptosis-associated speck-like protein containing a CARD, also called PYCARD) NEK7 (NIMA-related kinaseL7) to form the inflammasome complex^5, 6^. The NLRP3 inflammasome complex then recruits pro-Caspase-1 and promotes its auto-cleavage to form activated Caspase-1, which cleaves pro-IL-1β and pro-IL-18 to form mature IL-1β and IL-18^4,5^. Activated Caspase-1 also cleaves Gasdermin D (GSDMD)^7,8^. The released N-terminal half of GSDMD is inserted into plasma membrane and forms pores^9,10^ to release the mature IL-1β and IL-18^7,8,10^. Proteolytically activated GSDMD also leads to cell membrane rupture and cell death (pyroptosis)^8,9^, which contributes to the release of more intracellular components, including lactate dehydrogenase (LDH)^7,10^ and high mobility group box 1 (HMGB1)^11^.

NLRP3 is involved in many inflammatory and autoimmune diseases^4^. Gain-of-function NLRP3 mutations are relevant with cryopyrin-associated periodic syndromes (CAPS)^12–14^, a group of rare heritable autoinflammatory diseases that includes familial cold auto-inflammatory syndrome (FCAS), chronic infantile neurological, cutaneous, and articular (CINCA) syndrome, and Muckle–Wells syndrome (MWS)^15^. Hyperactivation of the NLRP3 inflammasome is associated with neurodegenerative diseases^16^, gout^17^, diabetes^18^, pathogen infection-induced septic shock^19,20^ and cytokine storm^21,22^. Therefore, accurate regulation of NLRP3 activation is critically important. Recent work shows that resting NLRP3 forms double-ring cages and is localized on *trans*-Golgi network (TGN). The TGN localization is crucial for NLRP3 activation and consequently ASC recruitment when responding to danger signals^23,24^. However, what regulates NLRP3 TGN localization is unknown. Understanding how NLRP3 localization is governed will provide new insight into the precise regulation of NLRP3 inflammasome, thus, providing potential targets for treating NLRP3-related human disorders.

*S*-palmitoylation is the addition of long-chain fatty acyl group, typically the 16-carbon palmitoyl group, to cysteine residues of target proteins via a thioester bond^25–27^. It is catalyzed by a group of palmitoyl acyltransferases, ZDHHCs (ZDHHC1-23 in human), which contain a conserved DHHC (Asp-His-His-Cys) cysteine-rich domain that is important for the enzymatic activity^25,28^. A few proteins involved in immune signaling have been identified to be modified by *S*-palmitoylation, such as STAT3^26^, NOD1/2^29,30^, STING^31^, and PD-L1^32^. Typically, protein *S*-palmitoylation promotes membrane targeting of otherwise soluble proteins through its hydrophobic chain^25,28,33^, which in turn affects protein-protein interaction and signal transduction^25,33^. Here, we report that NLRP3 can be *S*-palmitoylated by palmitoyl acyltransferase ZDHHC7 on Cys126 in macrophages, which was important for the TGN localization for resting NLRP3 and the dispersed TGN (dTGN) localization of activated NLRP3, ultimately promoting ASC recruitment and downstream NLRP3 inflammasome activation.

## Results

### NLRP3 is *S*-palmitoylated by ZDHHC7 in macrophages

Protein *S*-palmitoylation is considered to have important roles in inflammatory responses^25^. As NLRP3 is the key component in the NLRP3 inflammasome pathway and associated with many human diseases^4^, we hypothesized that it may be regulated by *S*-palmitoylation in macrophages. To investigate this possibility, we incubated the mouse bone marrow-derived macrophages (BMDMs) with a clickable palmitate analog, Alkyne 14 (Alk14), a chemical reporter for palmitoylation^26,34,35^, and performed click chemistry to conjugate Alk14-labeled proteins with biotin and determined the palmitoylation levels with streptavidin pull-down and immunoblot. The result showed NLRP3 was labeled with Alk14, suggesting that endogenous NLRP3 is palmitoylated BMDMs (Figure 1A). Similarly, the palmitoylation of NLRP3 was also detectable in phorbol 12-myristate 13-acetate (PMA)-primed human macrophage cell line, THP-1 (Figure 1B), suggesting NLRP3 palmitoylation occurred in both mouse and human macrophages.

**Figure 1.**
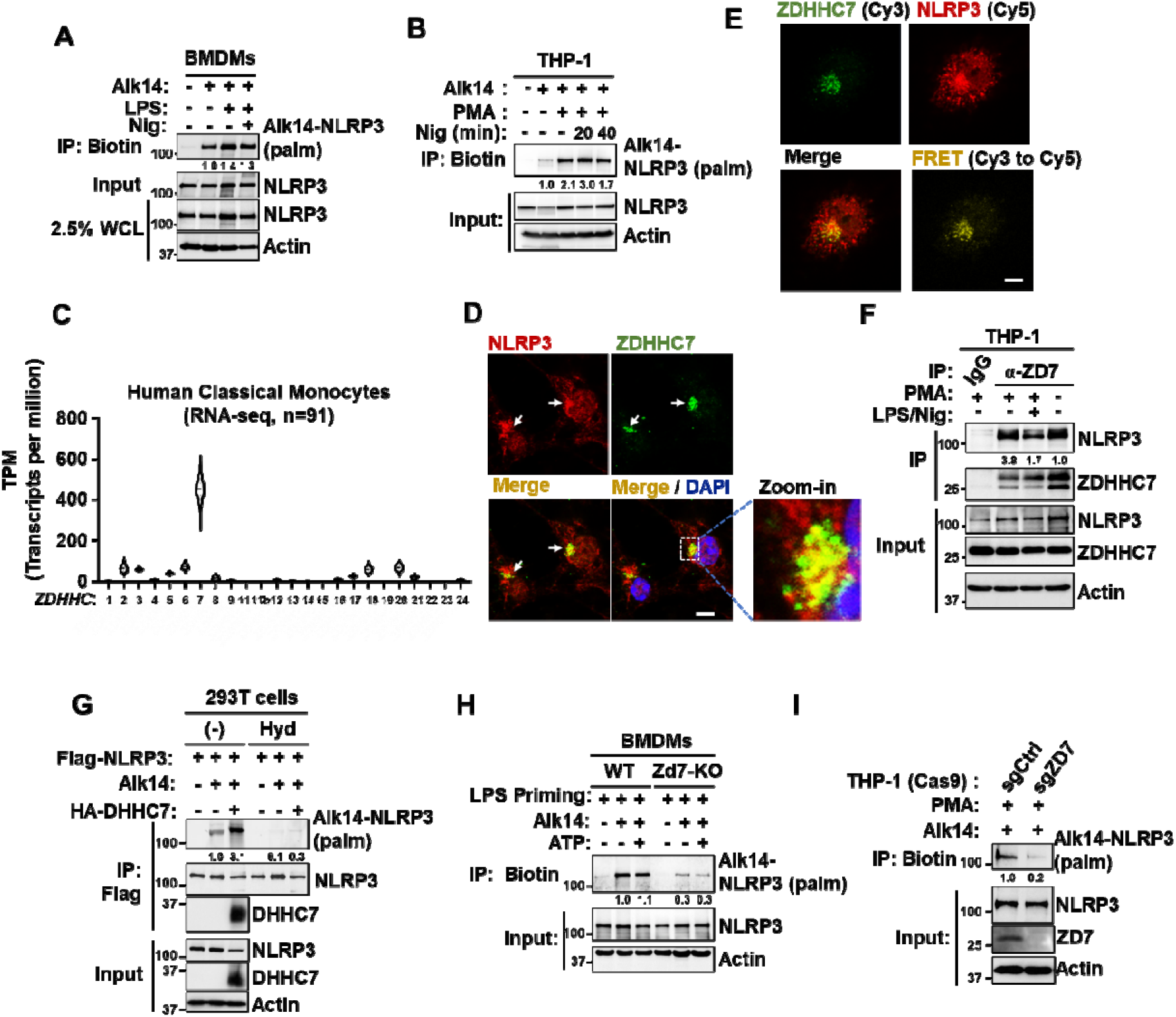
NLRP3 is *S*-palmitoylated by ZDHHC7 in macrophages. **(A)** Palmitoylation of NLRP3 in BMDMs by Alk14 labeling and click chemistry assay. BMDMs were incubated with 50 μM Alk14 for 6 h, during which LPS (200 ng/mL) was added and incubated for 4 h, then 10 μM of nigericin (Nig) wa added and incubated for 30 min. Cells were lysed and proteins were conjugated with biotin-azide, the labeled proteins were pulled down with streptavidin and blotted for NLRP3. NLRP3 palmitoylation level was quantified and normalized to NLRP3 protein levels in the input samples. **(B)** Palmitoylation of NLRP3 in PMA-primed THP-1 cells. Cells were primed with 10 ng/mL PMA for 24 h and then treated with Alk14, LPS, and nigericin as indicated, NLRP3 palmitoylation was determined as that in (**A**). **(C)** Expression of *ZDHHC* genes in human classical monocytes generated from the database of DICE^36^. (https://dice-database.org, n=91) **(D)** Representative confocal microscopic images of NLRP3 and ZDHHC7 in LPS-primed BMDMs. NLRP3 localization was detected with anti-NLRP3 staining, ZDHHC7 was detected with anti-ZDHHC7 staining, nucleus was stained with DAPI. Scale bar: 5 μm. **(E)** ZDHHC7 was stained with Cyanine 3 (Cy3) as a FRET donor and NLRP3 was stained with Cyanine 5 (Cy5) as a FRET acceptor in LPS-primed BMDMs, the interaction between NLRP3 and ZDHHC7 were determined by the FRET signal. Scale bar: 5 μm. **(F)** THP-1 cells were primed with or without PMA (10 ng/mL) for 24 h then treated with LPS and nigericin as indicated. Cell lysates were collected for immunoprecipitation (IP) and immunoblot analysis to detect the endogenous NLRP3-ZDHHC7 association. The pull downed NLRP3/ZDHHC7 ratio was calculated and normalized to protein levels in input samples. **(G)** Palmitoylation of Flag-NLRP3 in HEK 293T cells with ZDHHC7 expression was detected using Alk14 labeling and in-gel fluorescence. HEK 293T cells expressing Flag-NLRP3 were incubated with 50 μM of Alk14 for 6 h. Flag-NRLP3 was pulled down by IP and conjugated with TAMRA-azide using click chemistry. Palmitoylation of NLRP3 was detected using in-gel fluorescence after SDS-PAGE. Hydroxylamine treatment was used to confirm S-palmitoylation, which is hydroxylamine sensitive. NLRP3 palmitoylation level was quantified similarly as above. **(H)** Palmitoylation of NLRP3 is decreased by *Zdhhc7* KO in BMDMs. WT or *Zdhhc7*-KO (Zd7-KO) BMDMs were labeled with Alk14, primed with LPS, and activated with ATP as indicated, NLRP3 palmitoylation was determined as that in (**A**). **(I)** Palmitoylation of NLRP3 is decreased by *ZDHHC7* deletion in THP-1. NLRP3 palmitoylation level was quantified as above.

To identify which ZDHHC protein could possibly mediates NLRP3 modification, we determined the expression of all the *Zdhhc* genes (*Zdhhc1-9*, *Zdhhc11-25*) in BMDMs (Figure S1A) and analyzed the expression level of all the *ZDHHC* genes (*ZDHHC1-9, ZDHHC11, ZDHHC11b, ZDHHC12-24*) in human classical monocytes from public database^36^ (The Human Protein Atlas, n=91, Figure 1C). We found that several *Zdhhc* genes showed predominant expression level in BMDMs, including *Zdhhc7* (Figure S1A). In human classical monocytes, intriguingly, *ZDHHC7* showed a predominant expression compared with other *ZDHHC* genes (Figure 1C), suggesting *ZDHHC7* potentially has an important role in macrophages. Therefore, we investigated whether ZDHHC7 could mediate NLRP3 palmitoylation in macrophages.

We first determined whether ZDHHC7 interacts with NLRP3. Through immunofluorescence staining, we found NLRP3 co-localized with ZDHHC7 at resting state in LPS-primed BMDMs (Figure 1D). Fluorescence resonance energy transfer (FRET) assay further confirmed the strong interaction between NLRP3 and ZDHHC7 in LPS-primed BMDMs (Figure 1E). Additionally, co-immunoprecipitation (Co-IP) assay showed ZDHHC7 immunoprecipitation could pull down NLRP3 in THP-1 cells under resting, primed, or activated states (Figure 1F). The above data collectively suggested that NLRP3 associates with ZDHHC7 in macrophages.

We next co-expressed NLRP3 and ZDHHC7 in human embryonic kidney (HEK) 293T cells to see whether ZDHHC7 expression could increase NLRP3 palmitoylation. ZDHHC7 expression increased NLRP3 palmitoylation, and hydroxylamine treatment abolished ZDHHC7-mediated NLRP3 palmitoylation, suggesting the palmitoylation of NLRP3 is S-palmitoylation on cysteine residues (Figure 1G). Furthermore, deletion of *ZDHHC7* reduced the palmitoylation of NLRP3 expressed in HEK 293T cells (Figure S1B), and the enzymatically inactive mutant of ZDHHC7 (ZDHHS7) was unable to restore NLRP3 palmitoylation in *ZDHHC7*-deleted HEK 293T cells, in contrast to wildtype ZDHHC7 (Figure S1B), confirming that the enzymatic activity of ZDHHC7 was essential for NLRP3 palmitoylation. Lastly, to further confirm NLRP3 *S*-palmitoylation was induced by ZDHHC7 in macrophages, we isolated BMDMs from *Zdhhc7* knock out mouse, generated *ZDHHC7* knockout THP-1 cells and determined NLRP3 *S*-palmitoylation using Alk14 labeling. The results showed *Zdhhc7* deletion in mouse BMDMs, and *ZDHHC7* deletion in human THP-1 cells, dramatically diminished NLRP3 *S*-palmitoylation (Figure 1 H-I).

To investigate whether other ZDHHC proteins could mediate NLRP3 *S*-palmitoylation, we also expressed other ZDHHC in HEK 293T cells and performed NLRP3 Alk14 labeling. While several ZDHHC proteins (ZDHHC3, ZDHHC6, ZDHHC7, and ZDHHC9) could promote NLRP3 *S*-palmitoylation, ZDHHC7 exhibited the strongest activity (Figure S1C, D). To check if ZDHHC3, ZDHHC6, and ZDHHC9, which exhibited potential activity for NLRP3 *S*-palmitoylation in HEK 293T and high expression levels in BMDMs, could mediate NLRP3 *S*-palmitoylation in macrophages, we isolated BMDMs from *Zdhhc3*, *Zdhhc6*, or *Zdhhc9* knockout mouse and found that the knockout did not affect NLRP3 *S*-palmitoylation (Figure S1E, F). Taken together, the above data strongly demonstrated that NLRP3 is modified with *S*-palmitoylation mainly by ZDHHC7 in macrophages.

### ZDHHC7-catalyzed *S*-palmitoylation is important for NLRP3 inflammasome activation

We investigated if ZDHHC7-mediated *S*-palmitoylation regulates the process of inflammasome activation in macrophages. We first studied whether ZDHHC7 regulates the LPS priming process, especially the expression of NLRP3 inflammasome complex components. We found *Zdhhc7* KO did not affect LPS-induced p65 phosphorylation (Figure S2A), suggesting that ZDHHC7 did not regulate activation of NF-κB, the major transcription factor that mediates expression of inflammasome components during the LPS priming step. Consistent with this finding, *Zdhhc7* KO did not affect the LPS-induced mRNA expression of *Il-1b*/*Il-18* (Figure S2B) or protein levels of NLRP3, pro-Caspase-1, or full-length Gasdermin D (GSDMD) (Figure 2A), further validating that ZDHHC7 did not regulate the priming step. Next, to find out if ZDHHC7 affects the activation step of NLRP3 inflammasome, we determined the cleavage of Caspase-1 and GSDMD, and IL-1β secretion, which is the common readout of NLRP3 inflammasome activation. *Zdhhc7* KO potently inhibited Caspase-1 and GSDMD cleavage (Figure 2A), and IL-1β secretion (Figure 2B), after inflammasome activation in BMDMs, suggesting ZDHHC7 is important for the activation process of NLRP3 inflammasome in mouse macrophages. To verify this conclusion in human macrophages, we knocked out *ZDHHC7* in human THP-1 cells and found that *ZDHHC7* KO inhibited GSDMD cleavage and IL-1β secretion (Figure 2C, D). These data collectively indicated ZDHHC7 is important for NLRP3 inflammasome activation in mouse and human macrophages.

**Figure 2.**
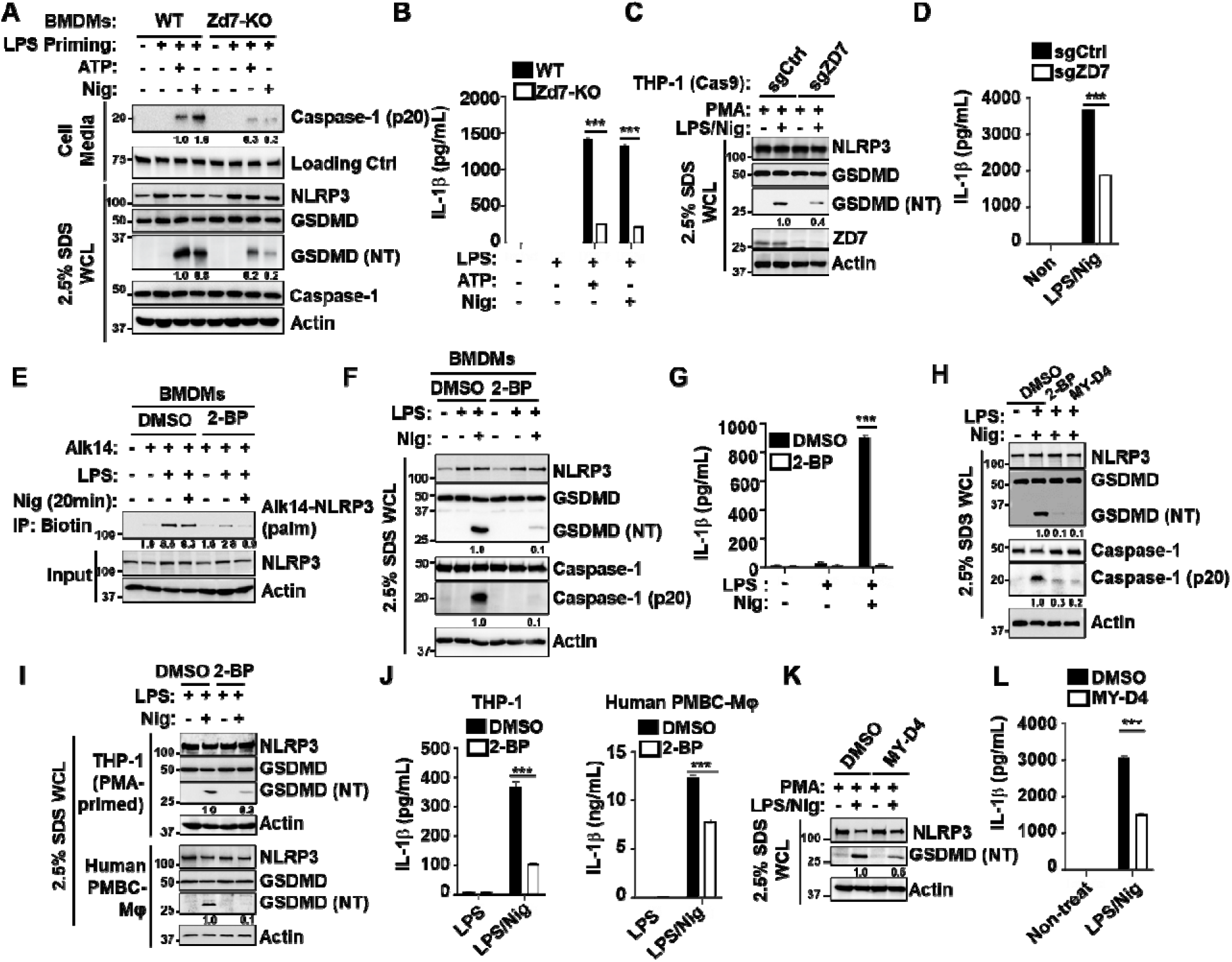
ZDHHC7 is important for NLRP3 inflammasome activation. **(A)** Immunoblot analysis of NLRP3, Caspase-1, GSDMD, and cleaved Caspase-1 (p20), cleaved GSDMD N-terminal domain (NT) in cell culture medium (top) and whole cell lysate (bottom) of WT and *Zdhhc7*-KO (Zd7-KO) BMDMs that were primed by LPS and activated with ATP or nigericin (Nig) as indicated. Cells were lysed with lysi buffer containing 2.5% SDS to get total proteins. **(B)** ELISA assay of IL-1β in cell culture media of WT and *Zdhhc7*-KO BMDMs. **(C)** Immunoblot analysis of NLRP3, GSDMD, and cleaved GSDMD N-terminal domain (NT) in *ZDHHC7*-depleted (sgZD7) and control (sgCtrl) THP-1 cells that were transduced with Cas9 nuclease. Cells were primed by PMA and activated with LPS and nigericin. Cells were lysed with lysis buffer containing 2.5% SDS to get total proteins. **(D)** IL-1β determined by ELISA in cell culture media of *ZDHHC7*-depleted (sgZD7) and control (sgCtrl) THP-1 cells that were primed with PMA and activated with LPS and nigericin. **(E)** *S*-Palmitoylation of NLRP3 is inhibited by 2-BP in BMDMs. BMDMs were pre-treated with 2-BP for 1 h before Alk14 incubation, and LPS and nigericin were added as indicated. NLRP3 palmitoylation level was quantified and normalized to NLRP3 protein level in input samples. **(F)** Immunoblot analysis of NLRP3, GSDMD, Caspase-1, cleaved Caspase-1 (p20), and cleaved GSDMD N-terminal domain (NT) in whole cell lysate of BMDMs treated with DMSO or 10 μM 2-BP and activated with 10 μM nigericin. **(G)** ELISA determination of IL-1β in cell culture media of LPS-primed BMDMs in (**F**). **(H)** Immunoblot analysis of NLRP3, GSDMD, Caspase-1, cleaved Caspase-1 (p20), and cleaved GSDMD N-terminal domain (NT) in whole cell lysate of BMDMs treated with DMSO, 10 μM 2-BP, or 20 μM MY-D4, and activated with 10 μM nigericin. **(I)** Immunoblot analysis of NLRP3, GSDMD, and cleaved GSDMD N-terminal domain (NT) in whole cell lysate of PMA-primed THP-1 (top) and human PBMC-derived primary macrophages (bottom) that were pre-treated with DMSO or 10 μM 2-BP for 1 h before activation with LPS and nigericin. **(J)** IL-1β determined by ELISA in cell culture media of THP-1 (left) and human PBMC-derived macrophages (right) in (**I**). **(K-L)** Immunoblot analysis of NLRP3 and cleaved GSDMD (NT) in whole cell lysate (**K**) and IL-1β determined by ELISA in cell culture media (**L**) of PMA-primed THP-1 cells pre-treated with DMSO or 20 μM MY-D4 for 1 h before activation with LPS and nigericin. Data with error bars are mean ± SEM. *p < 0.05, **p < 0.01, ***p < 0.001 as determined by unpaired Student’s t test.

To further confirm ZDHHC7-mediated *S*-palmitoylation is important for NLRP3 inflammasome activation, we studied NLRP3 inflammasome regulation by 2-bromopalmitate (2-BP), a pan-ZDHHC inhibitor, in BMDMs. 2-BP potently repressed NLRP3 *S*-palmitoylation in BMDMs (Figure 2E), but did not affect protein levels of NLRP3, pro-Capase-1, or GSDMD (Figure 2E, F), suggesting that 2-BP treatment did not regulate the priming step. 2-BP potently suppressed Caspase-1 and GSDMD cleavage, and IL-1β secretion in BMDMs activated by nigericin (Figure 2F, G), suggesting that 2-BP inhibited the activation step of NLRP3 inflammasome. As 2-BP was reported to have off-targets^37^, to further verify that effect of 2-BP in regulating NLRP3 inflammasome was through DHHC7, we synthesized a 2-BP-Alk probe that can be used to label proteins targeted by 2-BP (Figure S3A). Using 2-BP-Alk incubation, conjugation of biotin-azide and streptavidin pull-down, we verified that 2-BP-Alk could covalently bind to ZDHHC7 in inflammasome-activated BMDMs (Figure S3B). Importantly, 2-BP treatment did not further suppress GSDMD cleavage or pyroptosis (LDH release) in *Zdhhc7* knockout BMDMs activated with nigericin (Figure S3C, D), suggesting 2-BP inhibited NLRP3 inflammasome activation mainly through ZDHHC7. Additionally, MY-D4, a more recent improved inhibitor of ZDHHC enzymes^38^, also exhibited strong inhibition for Caspase-1 and GSDMD cleavage (Figure 2H) in BMDMs, further confirming ZDHHC inhibition could repress the NLRP3 inflammasome activation. We also evaluated the effect of ZDHHC inhibitors on inflammasome activation in human macrophages, including THP-1 and human peripheral blood monocyte cell (PBMC)-derived primary macrophages. 2-BP treatment inhibited GSDMD cleavage and IL-1β secretion in both PMA-differentiated THP-1 cells and human primary macrophages (Figure 2I, J). Another inhibitor MY-D4 also showed potent inhibition for GSDMD cleavage and IL-1β secretion in THP-1 (Figure 2K, L). The data collectively demonstrated that ZDHHC7-mediated *S*-palmitoylation is important for NLRP3 inflammasome activation in both mouse and human macrophages, which can be suppressed by ZDHHC inhibitors.

In addition to NLRP3 inflammasome, several other inflammasome complexes that respond to diverse danger signals have been studied in macrophages^39^. Non-canonical inflammasome is triggered by cytosolic LPS of invasive Gram-negative bacteria, which activates and oligomerizes Caspase-4/5/11 to cleave GSDMD and induce pyroptosis^40^. Absent in melanoma 2 (AIM2) senses double strand DNA from microbial pathogens or host cellular damage and recruits ASC to assemble the inflammasome complex, which activates Caspase-1 and leads to pyroptosis and cytokines secretion.^41–43^ To investigate whether ZDHHC7 regulates these NLRP3-independent inflammasome activation, we determined GSDMD cleavage and pyroptosis (LDH release) in WT and *Zdhhc7*-deleted BMDMs under non-canonical or AIM2-mediated inflammasome conditions. We found that *Zdhhc7* deletion did not affect non-canonical-inflammasome-induced GSDMD cleavage or pyroptosis in BMDMs (Figure S4A, B). Zdhhc7 deletion also did not affect GSDMD cleavage induced by AIM2 inflammasome (Figure S4C). Thus, ZDHHC7 does not regulate non-canonical or AIM2 inflammasomes activation.

Lastly, to check if ZDHHC3, ZDHHC6, or ZDHHC9, which exhibited high expression in BMDMs, may regulate NLRP3 activation in macrophages, we also isolated BMDMs from *Zdhhc3*, *Zdhhc6*, or *Zdhhc9* knock out mouse, and found that none of them affected GSDMD cleavage indued by ATP or nigericin (Figure S5A-C), suggesting that these ZDHHC proteins are not involved in NLRP3 activation.

### NLRP3 was *S*-palmitoylated by ZDHHC7 on Cys126

Thus far, our data demonstrated that ZDHHC7 can palmitoylate NLRP3 and promote NLRP3 inflammasome activation. However, whether the effect of ZDHHC7 on NLRP3 inflammasome activation is through NLRP3 *S*-palmitoylation or through other substrate proteins was not clear. To address this, we sought to identify the modified cysteine(s) on NLRP3 by ZDHHC7. Once the modified cysteine is identified, the cysteine mutant can be used further validate that ZDHHC7’s effect on NLRP3 inflammasome activation is through NLRP3 palmitoylation.

Recent work reported that resting state NLRP3 localizes to the TGN by ionic bonding between its conserved polybasic region and the negatively charged phosphatidylinositol-4-phosphates (PtdIns4P) on the TGN^23,24,44^. We hypothesize that ZDHHC7 may palmitoylate NLRP3 and promote the TGN localization. We screened the cysteine residues on NLRP3 and found there are two cysteines, Cys126 and Cys133, that are near the polybasic region (Figure 3A). Cys126 (human Cys130) is conserved in all NLRP3 examined while Cys133 is not. We mutated each of them to serine and performed Alk14 labeling. Mutating Cys126 to serine (C126S), but not Cys133 to serine (C133S), dramatically decreased the S-palmitoylation level of NLRP3 (Figure 3B). Additionally, C126S mutant of mouse NLRP3 or C130S mutant of human NLRP3 dramatically decreased ZDHHC7-promoted NLRP3 *S*-palmitoylation in HEK 293T cells (Figure 3C, D), suggesting Cys126 (human Cys130) is the major site of NLRP3 S-palmitoylation mediated by ZDHHC7.

**Figure 3.**
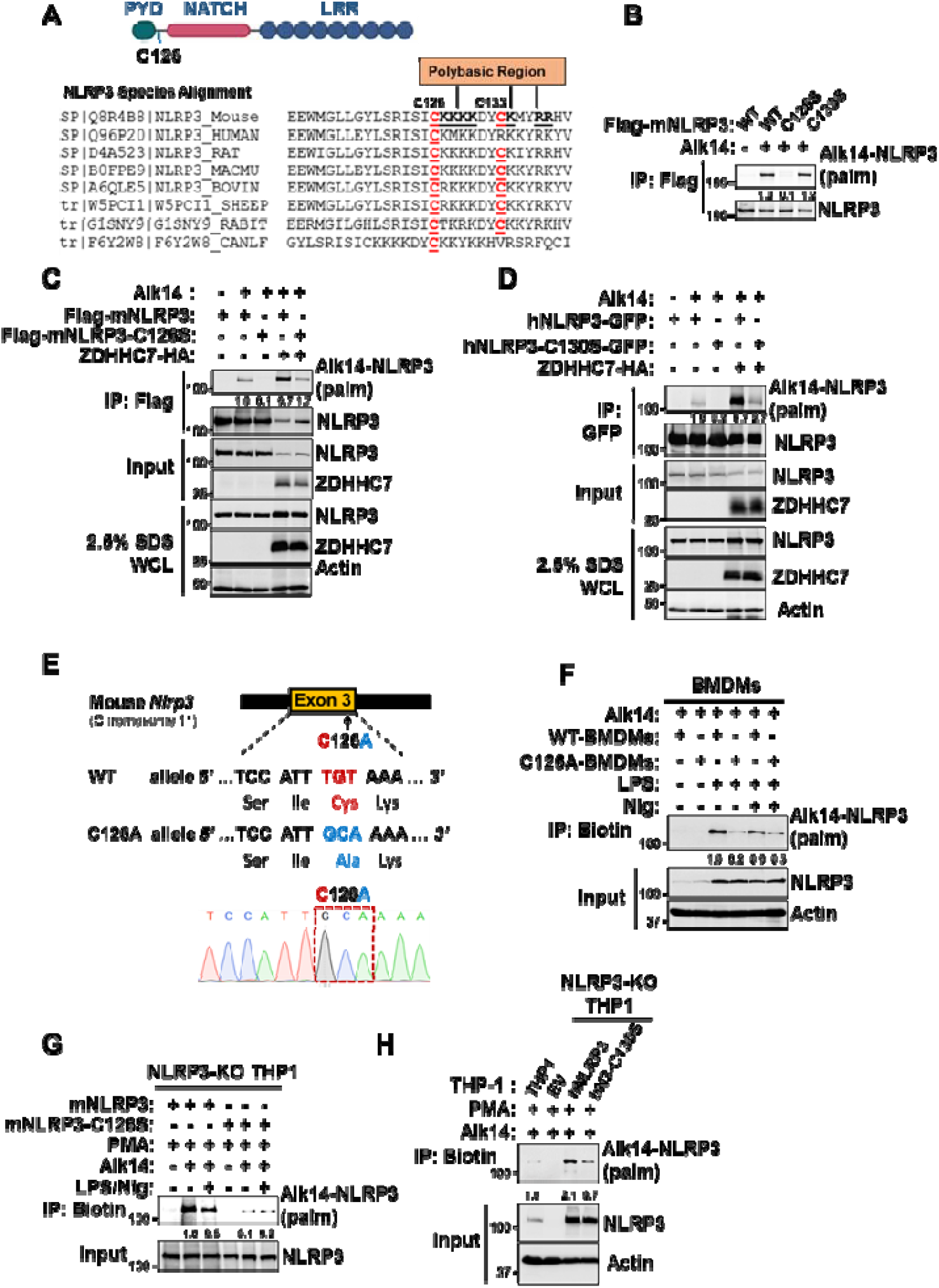
NLRP3 was *S*-palmitoylated by ZDHHC7 on Cys126. **(A)** Schematic diagram of NLRP3 domains showing the location of Cys126 (top), and amino acid sequence alignment of NLRP3 from different species highlighting the conserved Cys126 and the nearby Cys133 in red (bottom). The TGN binding polybasic region is shown in bold and underlined. **(B)** Palmitoylation of mouse NLRP3 WT and cysteine to serine mutants (CS) in HEK 293T cells using Alk14 labeling, click chemistry assay with TAMRA-azide, and in-gel fluorescence analysis. NLRP3 palmitoylation level was quantified and normalized to NLRP3 protein levels in the input samples. **(C-D)** Palmitoylation of mouse (**C**) or human (**D**) NLRP3 WT and cysteine to serine mutant (C126S of mouse NLRP3, C130S of human NLRP3) in HEK293T cells. NLRP3 S-palmitoylation were determined using Alk14 labeling as in (**B**). C126S or C130S mutant dramatically decreased basal level or ZDHHC7 expression-promoted NLRP3 palmitoylation. NLRP3 palmitoylation level was quantified as above. **(E)** The generation of the CRISPR-edited NLRP3 C126A mutant (upper panel) and sequencing verification of the codon replacement (lower panel). **(F)** Palmitoylation of NLRP3 in WT and C126A BMDMs by Alk14 labeling. BMDMs were incubated with Alk14, LPS and nigericin (Nig) as indicated. NLRP3 palmitoylation level was quantified as above. **(G)** Palmitoylation of mouse NLRP3 WT and C126S mutant that were reconstituted stably in NLRP3-deleted THP-1 cells. Cells were differentiated with PMA for 24 h and labeled with 50 μM Alk14 for 6 h, during which LPS was treated for 4 h before nigericin activation for another 1 h. After cell lysis, the labeled proteins were conjugated with biotin-azide, pulled down with streptavidin resin, and blotted for NLRP3. NLRP3 palmitoylation level was quantified as above. **(H)** Palmitoylation of human NLRP3 WT and C130S mutant that were reconstituted stably in NLRP3-deleted THP-1 cells. Cells were differentiated with PMA for 24 h and labeled with 50 μM Alk14 for 6 h, the labeled proteins were detected as in (**G**).

To verify that NLRP3 Cys126 is the major physiological *S*-palmitoylation site in macrophages, we generated C126A *Nlrp3* gene-edited mouse, with the *TGT of Nlrp3* alleles in Exon 3, encoding Cys126, mutated to alanine-encoded sequence *GCA* (*TGT*>*GCA*, Figure 3E), and determined NLRP3 S-palmitoylation in BMDMs. We found C126A mutant dramatically diminished the S-palmitoylation of NLRP3 in both LPS-primed and nigericin-activated BMDMs (Figure 3F), confirming that Cys126 is the major *S*-palmitoylation site in mouse primary macrophages. To investigate whether the corresponding Cys130 in human NLRP3 is the major palmitoylation site in human macrophages, we depleted endogenous NLRP3 in THP-1 cells and then reconstituted with either the wildtype or cysteine mutant NLRP3. Both mouse C126S and human C130S NLRP3 mutant had dramatically decreased modification level in THP-1 cells compared to wildtype NLRP3 (Figure 3G, H). Therefore, the above data collectively indicates Cys126 is the major site of NLRP3 *S*-palmitoylation mediated by ZDHHC7 in macrophages.

### NLRP3 Cys126 palmitoylation by ZDHHC7 is crucial for NLRP3 inflammasome activation in macrophages

To confirm the importance of Cys126 palmitoylation for NLRP3 inflammasome activation, we determined NLRP3 inflammasome activation in BMDMs differentiated from wildtype, CRISPR-edited C126A, and *Nlrp3*-KO mouse. We found Caspase-1 and GSDMD cleavage (Figure 4A, B), as well as IL-1β and IL-18 secretion in C126A BMDMs was dramatically inhibited compared with that in wildtype BMDMs (Figure 4C, D). The effect of C126A mutation was close to that in *Nlrp3*-KO BMDMs (Figure 4A-D). Furthermore, we also compared the effect of *Zdhhc7* knockdown on inflammasome activation with NLRP3 WT and C126A mutant in BMDMs. *Zdhhc7* knockdown suppressed Caspase-1 and GSDMD cleavage in NLRP3 wildtype BMDMs, but not the NLRP3 C126A mutant BMDMs (Figure 4E, Figure S6A), supporting that ZDHHC7’s effect on NLRP3 inflammasome activation is mainly through C126 palmitoylation. Similarly, 2-BP treatment significantly decreased Caspase-1 and GSDMD cleavage in wildtype BMDMs but not in C126A mutant BMDMs (Figure 4F). Therefore, Cys126 palmitoylation is crucial for NLRP3 inflammasome activation in mouse primary macrophages and the effect of ZDHHC7 on NLRP3 inflammasome activation is mainly through C126 palmitoylation.

**Figure 4.**
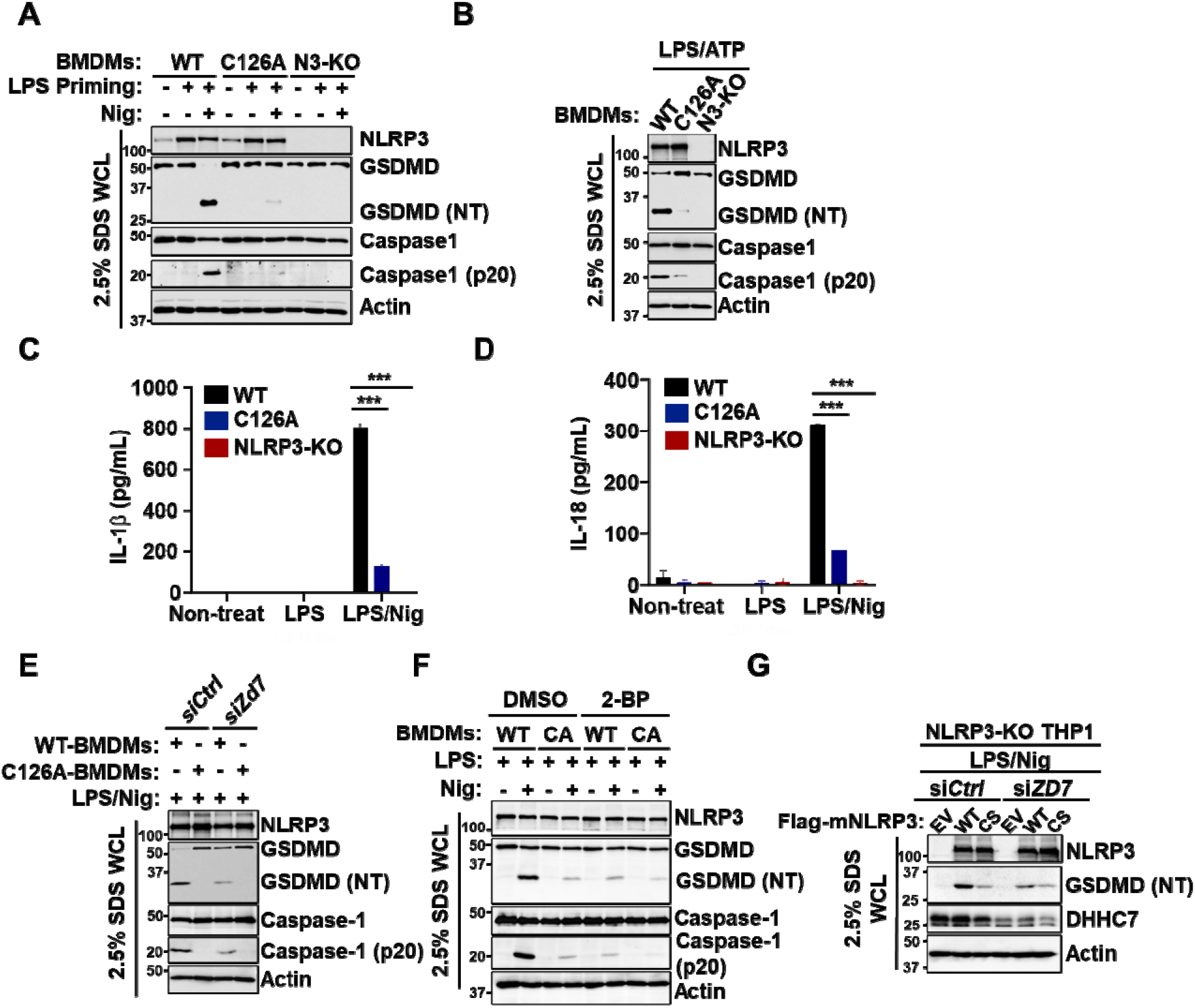
NLRP3 Cys126 mutant inhibited NLRP3 inflammasome activation in macrophages. **(A-B)** Immunoblot analysis of NLRP3, Caspase-1, GSDMD, cleaved Caspase-1 (p20), and cleaved GSDMD (NT) in the whole cell lysate of NLRP3 WT, C126A, and KO (N3-KO) BMDMs that were primed with LPS and activated with nigericin (Nig, **A**) or ATP (**B**). Cells were lysed with lysis buffer containing 2.5% SDS to get total proteins. **(C-D)** Levels of IL-1β (**C**) and IL-18 (**D**) in the cell culture media of LPS-primed BMDMs in (**A**). **(E)** Immunoblot analysis of NLRP3, Caspase-1, GSDMD, cleaved Caspase-1 (p20), and cleaved GSDMD (NT) in the whole cell lysate of NLRP3 WT and C126A mutant BMDMs that were *Zdhhc7* knockdown (si*Zd7*), primed by LPS, and activated with nigericin. **(F)** Immunoblot analysi of NLRP3, Caspase-1, GSDMD, cleaved Caspase-1 (p20), and cleaved GSDMD (NT) in the whole cell lysate of NLRP3 WT and C126A mutant BMDMs that were primed with LPS and treated with DMSO or 10 μM 2-BP before activation with nigericin as indicated. **(G)** Immunoblot analysis of NLRP3, ZDHHC7, and GSDMD cleavage (NT) in NLRP3-deleted THP-1 cells that were reconstituted with WT or C126S mutant NLRP3 and deleted of *ZDHHC7* (si*ZD7*). Cells were PMA-differentiated, LPS primed, and stimulated with 10 μM nigericin for 1h as indicated. Data with error bars are mean ± SEM. *p < 0.05, **p < 0.01, ***p < 0.001 as determined by the unpaired Student’s t test.

We also investigate whether NLRP3 Cys126 palmitoylation also had a vital function in human macrophages. We introduced wildtype and C126S NLRP3 in THP-1 cells where the endogenous NLRP3 is deleted. Wildtype NLRP3 reconstitution restored GSDMD cleavage and IL-1β secretion, but C126S reconstitution did not (Figure S6B-D). Moreover, *ZDHHC7* knockdown suppressed GSDMD cleavage induced by wildtype NLRP3, but not by C126S mutant (Figure 4G), suggesting the effect of ZDHHC7 on NLRP3 inflammasome activation is mainly through C126 palmitoylation. Similarly, 2-BP treatment decreased GSDMD cleavage and IL-1β secretion induced by wildtype NLRP3 but not by C126S mutant in the THP-1 cells that were depleted of endogenous NLRP3 (Figure S6E, F). Taken together, these data support that ZDHHC7-mediated NLRP3 Cys126 *S*-palmitoylation is important for NLRP3 inflammasome activation in human macrophages.

### ZDHHC7-mediated Cys126 palmitoylation regulates NLRP3 inflammasome activation *in vivo*

Next, we sought to test whether ZDHHC7-mediated NLRP3 Cys126 palmitoylation is also functioning *in vivo*. The LPS intraperitoneal injection-induced endotoxic shock and mono sodium urate crystal (MSU)-induced peritonitis mouse models are widely used to assess NLRP3-dependent inflammasome responses *in vivo* ^45–48^. Therefore, we used these two models in wildtype and *Zdhhc7* KO mice to validate the regulation of NLRP3 *S*-palmitoylation *in vivo*. *Zdhhc7* knock-out mice had lower IL-1β and IL-18 secretion (Figure 5A), while TNFα secretion, which is mediated by NF-κB signaling, was similar in *Zdhhc7* KO and wildtype mice (Figure 5B), suggesting that *Zdhhc7* KO inhibits inflammasome activation *in vivo* but not NF-κB activation. Similarly, 2-BP administration in mice also inhibited LPS-induced IL-1β and IL-18 secretion (Figure 5C), but not TNFα or IL-6 secretion (Figure 5D). Furthermore, 2-BP administration alleviated the mortality of endotoxic shock animals (Figure 5E). More importantly, CRISPR-edited NLRP3 C126A also protected mice from endotoxic shock, with lower circulating IL-1β and HMGB1 in NLRP3 C126A mice (Figure 5F). Additionally, *Zdhhc7* KO mice were more resistant to MSU-induced peritonitis, with lower neutrophils and macrophages infiltration in peritoneal lavage fluids and diminished IL-1β level in the serum of *Zdhhc7* KO mice (Figure 5G, H). All the above data were consistent with the cellular results and confirmed that NLRP3 *S*-palmitoylation regulated NLRP3 inflammasome activation i*n vivo*.

**Figure 5.**
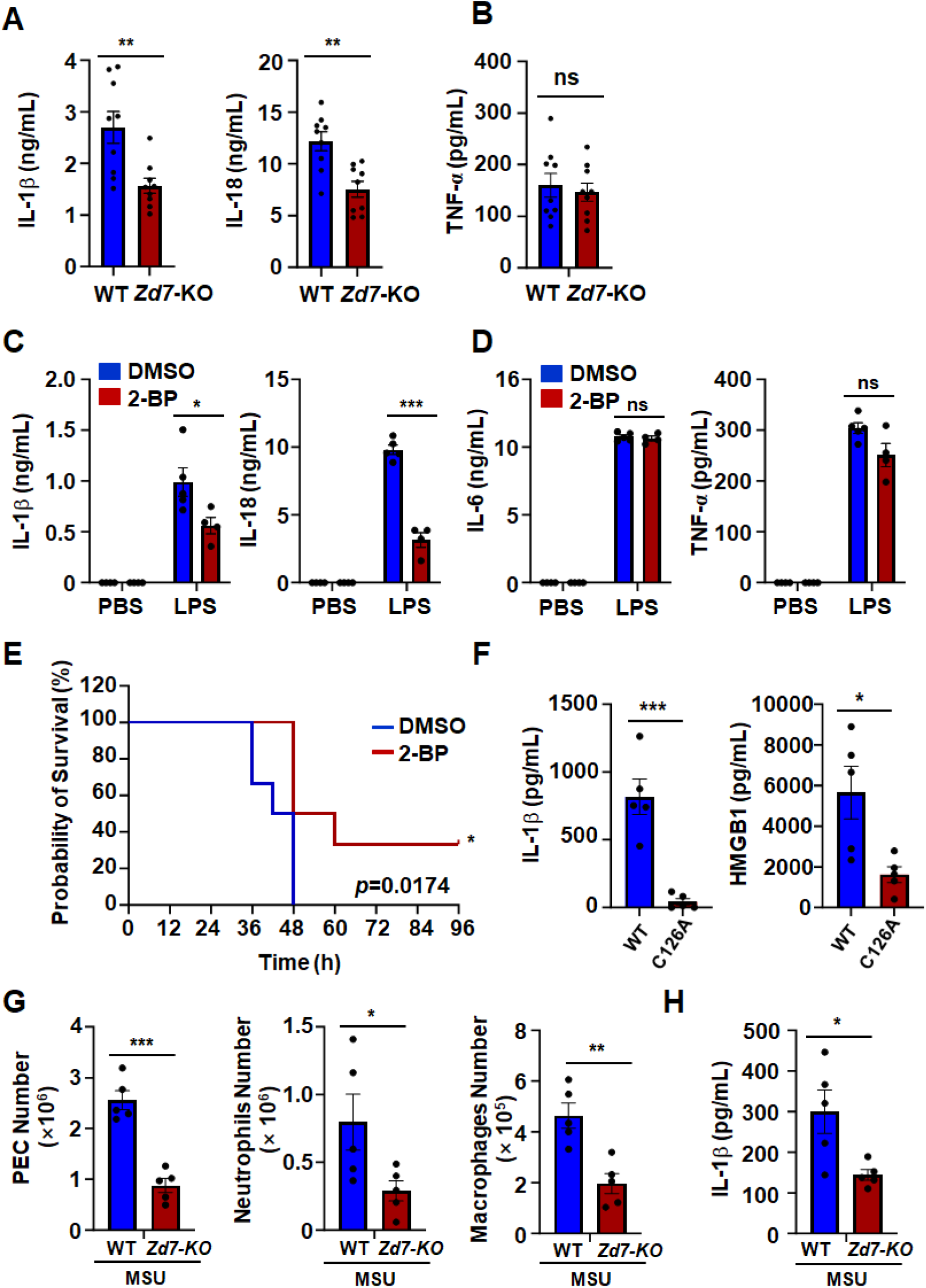
ZDDHC7-promoted Cys126 palmitoylation affects NLRP3 inflammasome activation *in vivo*. **(A-B)** Mouse serum IL-1β and IL-18 (**A**), and TNF-α (**B**) measured by ELISA for ZDHHC7 WT and KO (*Zd7*-KO) mice with LPS-induced endotoxic shock (n = 9 for each group). Mice were administered with LPS (35 mg/kg) by intraperitoneal injection and 8 h later, mouse serum was collected for ELISA determination. **(C-D)** Mouse serum IL-1β and IL-18 (**C**), and IL-6 and TNF-α (**D**) measured by ELISA for WT mice with endotoxic shock after administration of DMSO or 25 mg/kg 2-BP (LPS: n = 5 or 4; PBS: n = 4, for each group). **(E)** Survival data of WT mice (n = 6 for each group) in response to LPS (35 mg/kg, i.p.) challenge after administration of DMSO or 25 mg/kg 2-BP. **(F)** Mouse serum IL-1β and HMGB1 measured by ELISA for NLRP3 WT and C126A mice with endotoxic shock (n = 5 for each group). **(G-H)** Numbers of peritoneal exudate cells (PEC), infiltrated neutrophils and macrophages in peritoneal lavage fluids (**G**) and ELISA of IL-1β in the serum (**H**) of ZDHHC7 WT and KO (*Zd7*-KO) mice that were intraperitoneally injected (i.p.) with LPS (1 mg/kg mice weight, in PBS) for 3 h and then 200 μl of MSU (10 mg/ml, in PBS) for 6□h (n = 5, for each group). Data with error bars are mean ± SEM. *p < 0.05, **p < 0.01, ***p < 0.001 as determined by unpaired Student’s t test.

### ZDHHC7-catalyzed Cys126 palmitoylation promotes NLRP3 *trans*-Golgi network localization

We next investigated how Cys126 palmitoylation promotes NLRP3 inflammasome activation. *S*-palmitoylation promotes protein anchoring into the membrane lipid bilayer, which in turn regulates protein subcellular localization and protein-protein interactions^26,29,31^. For instance, ZDHHC7-catalyzed STAT3 *S*-palmitoylation mediates its plasma membrane localization and JAK2-STAT3 interaction, promoting STAT3 phosphorylation and signaling transduction^26^. Recent work elucidated that resting NLRP3 is mainly membrane-bound and forms oligomeric double-ring cages on the TGN, which is necessary for NLRP3 inflammasome activation^23,24,49^. Therefore, we investigated whether NLRP3 *S*-palmitoylation regulates its TGN localization. By expressing NLRP3 fused with GFP in HEK 293 T cells, we found that resting NLRP3 indeed co-localized with the TGN markers, 58K Golgi (Figure 6A, B) or TGN38 (Figure 6C, Figure S7A). Furthermore, 2-BP treatment, or NLRP3 *S*-palmitoylation site mutants, or *ZDHHC7* deletion significantly inhibited resting NLRP3 localization on the TGN (Figure 6A, B, Figure S7A). Therefore, NLRP3 Cys126 *S*-palmitoylation was important for resting NLRP3 TGN localization.

**Figure 6.**
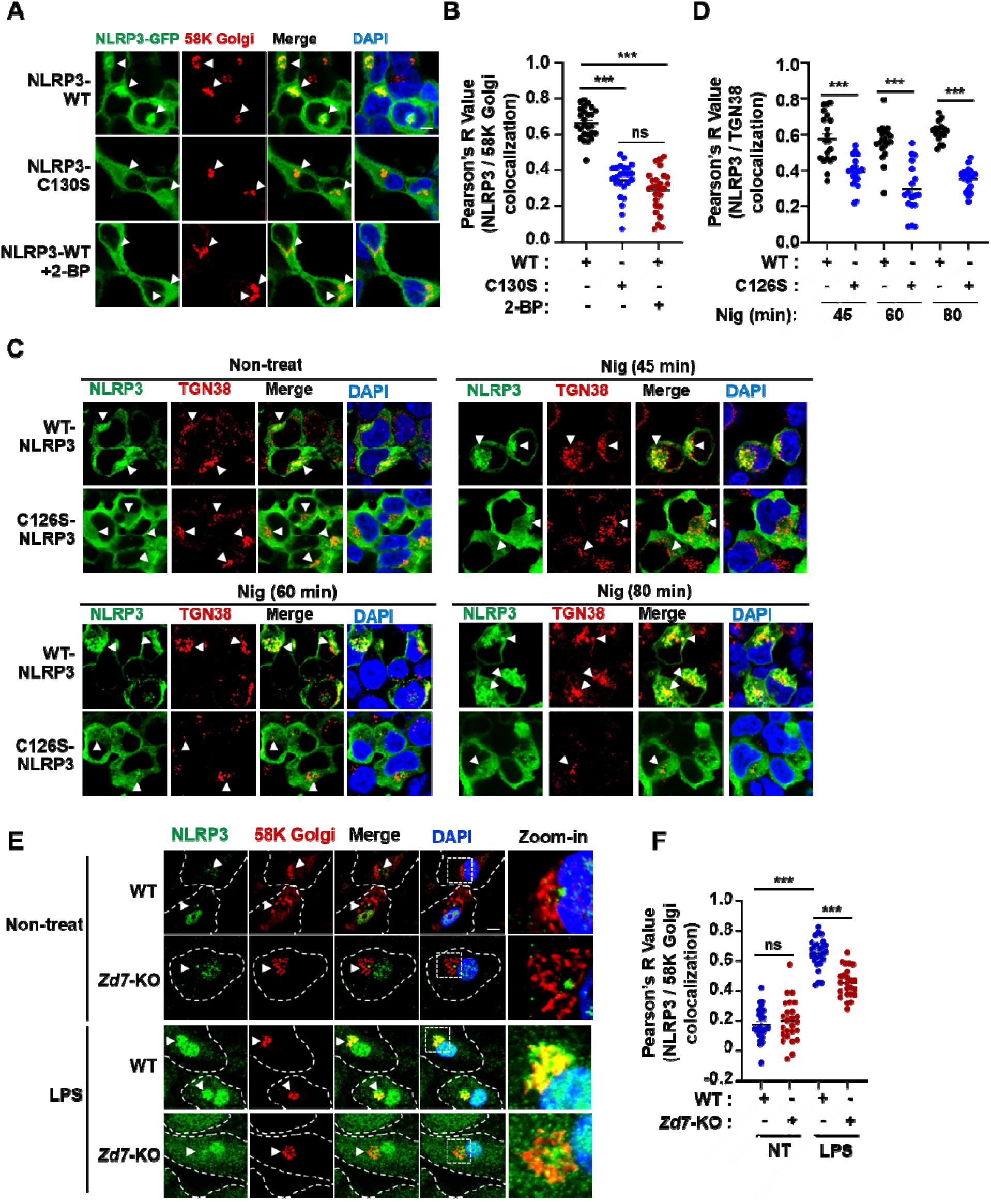
NLRP3 Cys126 palmitoylation promotes NLRP3 localization on *trans*-Golgi network. **(A)** Representative confocal microscopic images of GFP-tagged human NLRP3 WT or C130S mutant expressed in HEK293T cells treated with DMSO or 50 μM 2-BP for 12 hours. NLRP3-GFP was shown in green, TGN localization was stained with anti-58K Golgi (red), nucleus was stained with DAPI (blue). Scale bar: 5 μm. **(B)** Quantification of images in (**A**) by Pearson’s correlation coefficient using ImageJ, ranging from −1 to 1. **(C)** Confocal microscopic images of WT and C126S mutant of mouse NLRP3-Flag in HEK 293T cells that were treated with or without nigericin (Nig) for indicated times. NLRP3 was stained with anti-Flag (green), TGN was stained with anti-TGN38 (red), and nucleus was stained with DAPI (blue). WT but not the C126S mutant of NLRP3 colocalized with the TGN marker, TGN38. Scale bar: 5 μm. **(D)** Quantification of images in (**C**) by Pearson’s correlation coefficient using ImageJ, ranging from -1 to 1. **(E)** Representative confocal microscopic images for endogenous NLRP3 in non-treated or LPS-primed WT and *Zdhhc7* KO (*Zd7*-KO) BMDMs. NLRP3 localization was detected with anti-NLRP3 staining, TGN localization was stained with anti-58K Golgi, nucleus was stained with DAPI. Scale bar: 5 μm. **(F)** Quantification of images in (**E**) by Pearson’s correlation coefficient using ImageJ. Data with error bars are mean ± SEM. *p < 0.05, **p < 0.01, ***p < 0.001 as determined by unpaired Student’s t test.

After NLRP3 is activated, TGN will be disrupted to form the dispersed TGN (dTGN) vesicle, which will carry NLRP3 to microtubular organization center (MTOC)^46^, where NLRP3 will recruit NEK7 and the adaptor protein ASC to form the inflammasome complex^46,50^. Our data showed that nigericin-activated NLRP3 was localized on dispersed TGN (Figure 6C), which was consistent with previous reports^23,46^, whereas C126S mutant abolished the NLRP3 dTGN localization (Figure 6C, D), indicating that NLRP3 Cys126 *S*-palmitoylation was also important for its dTGN localization after NLRP3 activation. Endogenous NLRP3 was localized on TGN at LPS-primed resting state, while *Zdhhc7* KO inhibited the endogenous NLRP3 TGN localization in BMDMs (Figure 6E, F). Interestingly, Golgi-localized ZDHHC7 was dispersed on the dTGN during inflammasome activation (Figure S7B), and NLRP3 continued to be co-localized with ZDHHC7 after NLRP3 activation in BMDMs (Figure S7C). Therefore, the above data strongly suggested that ZDHHC7-mediated NLRP3 Cys126 *S*-palmitoylation was necessary for NLRP3 TGN/dTGN localization.

### ZDHHC7-catalyzed Cys126 palmitoylation promotes ASC recruitment and inflammasome assembly after NLRP3 activation

NLRP3 dTGN localization acts as an oligomerization scaffold for downstream adaptor ASC recruitment after inflammasome activation and trafficking to MTOC^23,24,51^. Thus, we studied whether ZDHHC7-mediated Cys126 palmitoylation would affect the recruitment of ASC. We performed Co-IP assays by expressing NLRP3 and ASC proteins in HEK 293T cells and found that 2-BP treatment inhibited NLRP3 interaction with ASC in HEK 293T cells (Figure 7A). NLRP3 C126S mutation also decreased the interaction of NLRP3 and ASC, while 2-BP treatment had no effect on the interaction between ASC and NLRP3 C126S (Figure 7A), suggesting that the effect of 2-BP on NLRP3-ASC interaction is mainly through inhibiting NLRP3 Cys126 palmitoylation. As expected, *ZDHHC7* KO in HEK 293T cells strongly suppressed NLRP3 WT interaction with ASC (Figure 7B) but had little effect on NLRP3 C126S mutant interaction with ASC, suggesting that ZDHHC7-mediated Cys126 palmitoylation contributed to the recruitment of ASC. To further confirm this in a more physiological setting, we examined the effect of *Zdhhc7* KO, 2-BP treatment, or NLRP3 C126A mutation on the interaction of NLRP3 with ASC after inflammasome activation in BMDMs. Both *Zdhhc7* KO and 2-BP decreased the interaction of NLRP3 with ASC (Figure 7C, D), and NLRP3 C126A mutant had a dramatically decreased interaction with ASC in BMDMs compared to NLRP3 wildtype (Figure 7E). The data collectively demonstrated that ZDHHC7-promoted NLRP3 Cys126 palmitoylation was critical for ASC recruitment after NLRP3 activation in macrophages.

**Figure 7.**
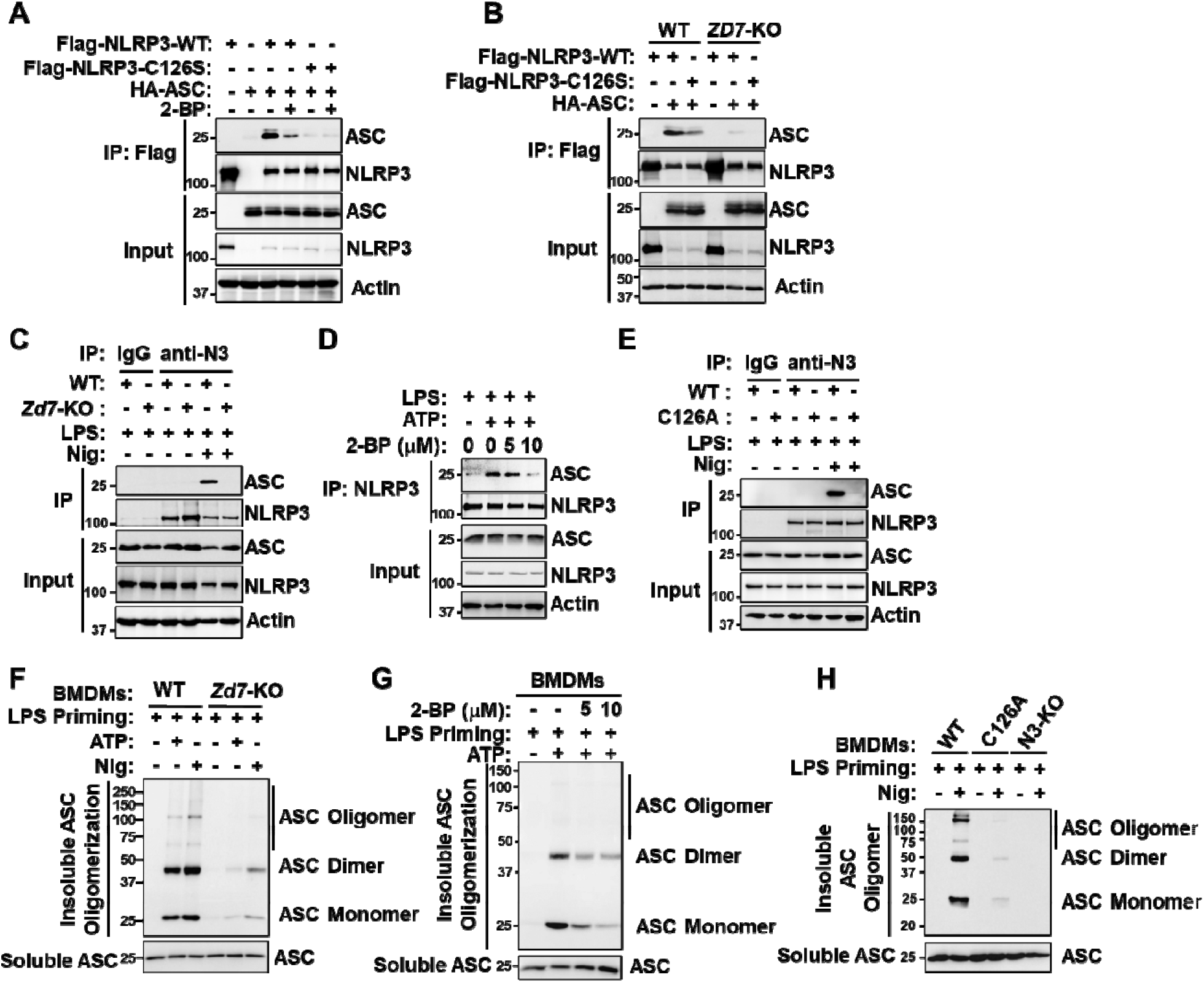
Cys126 palmitoylation promotes ASC recruitment and oligomerization after inflammasome activation. **(A)** Immunoblot analysis of NLRP3 interaction with ASC expressed in HEK 293T cells by co-IP with DMSO or 50 μM 2-BP treatment for 12 h. Interaction of ASC with WT NLRP3 was decreased by 2-BP, interaction of ASC with NLRP3 C126S mutant was weaker than that with WT NLRP3 and was not further affected by 2-BP. **(B)** Immunoblot analysis of NLRP3 interaction with ASC expressed in WT or *ZDHHC7*-KO (*ZD7*-KO) HEK 293T cells by co-IP. The interaction of ASC with NLRP3 C126S mutant was weaker than that with WT NLRP3 and ZDHHC7 KO dramatically decreased the interaction of ASC with WT NLRP3. **(C)** NLRP3 interaction with ASC in WT or *Zdhhc7*-KO BMDMs. Cells were primed with 200 ng/ml LPS for 4 h and activated with 10 μM nigericin (Nig) for 45 min. IP was done with anti-NLRP3 or isotype IgG control and immunoblot was done with anti-ASC and anti-NLRP3. The interaction of ASC with NLRP3 after inflammasome activation in *Zdhhc7*-KO BMDMs was almost abolished when compared with that in WT BMDMs. **(D)** Immunoblot analysis of NLRP3 interaction with ASC in mouse macrophage cell line J774A.1 treated with DMSO or 2-BP. The cells were primed with 200 ng/mL LPS for 4 h, then incubated with DMSO or 2-BP (5 and 10 μM) and ATP (5 mM) for 30 min. **(E)** NLRP3 interaction with ASC in WT or *Nlrp3* C126A BMDMs. Cells were primed with LPS and activated with nigericin. The interaction of ASC with NLRP3 after inflammasome activation in C126A BMDMs was almost abolished when compared to that in WT BMDMs. **(F)** Immunoblot analysi of insoluble ASC and its oligomerization. Disuccinimidyl suberate (DSS) was used to cross-link ASC in LPS-primed ZDHHC7 WT or KO BMDMs activated with ATP or nigericin. **(G)** Immunoblot analysis of insoluble ASC and its oligomerization (detected after DSS cross-linking) in LPS-primed WT BMDMs that were treated with DMSO or 2-BP (5 or 10 μM) and activated with ATP. **(H)** Immunoblot analysis of insoluble ASC and its oligomerization (detected after DSS cross-linking) in LPS-primed NLRP3 WT, C126A, or KO BMDMs that were activated with nigericin.

As ASC recruitment is important for the inflammasome complex assembly, we evaluated the regulation of NLRP3 Cys126 palmitoylation on inflammasome complex formation, especially the ASC oligomerization. Our data indicated that *Zdhhc7* knockout, 2-BP treatment, or NLRP3 C126A mutation dramatically inhibited ASC oligomerization induced by NLRP3 activation (Figure 7F-H), supporting that ZDHHC7-mediated NLRP3 Cys126 palmitoylation is important for inflammasome complex assembly after NLRP3 activation. Similarly, we found ASC oligomerization could be rescued by wildtype NLRP3 reconstitution in NLRP3 KO human THP-1 cells, but not by NLRP3 C126S mutant reconstitution (Figure S8). Therefore, ZDHHC7-catalyzed Cys126 palmitoylation is important for ASC recruitment and inflammasome assembly after NLRP3 activation in macrophages.

### ZDHHC7-mediated NLRP3 activation is different from ZDHHC12-mediated NLRP3 suppression

Thus far, our results strongly suggest that ZDHHC7-catalyzed NLRP3 Cys126 S-palmitoylation is important for NLRP3 inflammasome activation through regulating NLRP3 TGN/dTGN localization and ASC recruitment in macrophages. However, recent work reported that NLRP3 was palmitoylated on Cys844 by ZDHHC12 and this negatively regulated NLRP3 inflammasome activation^52^. Therefore, we compared ZDHHC7-catalyzed Cys126 palmitoylation and ZDHHC12-catalyzed Cys841 modification. We deleted *ZDHHC7* or *ZDHHC12* in THP-1 cells to assess the NLRP3 *S*-palmitoylation. *ZDHHC7* deletion decreased the *S*-palmitoylation of NLRP3 in both the PMA-primed and inflammasome-activated THP-1 cells, whereas *ZDHHC12* deletion only decreased the NLRP3 *S*-palmitoylation in inflammasome-activated THP-1 cells (Figure 8A). This suggested that NLRP3 *S*-palmitoylation mediated by ZDHHC7 can occur regardless of the activation stage, while ZDHHC12-catalyzed NLRP3 *S*-palmitoylation was induced after NLRP3 activation.

**Figure 8.**
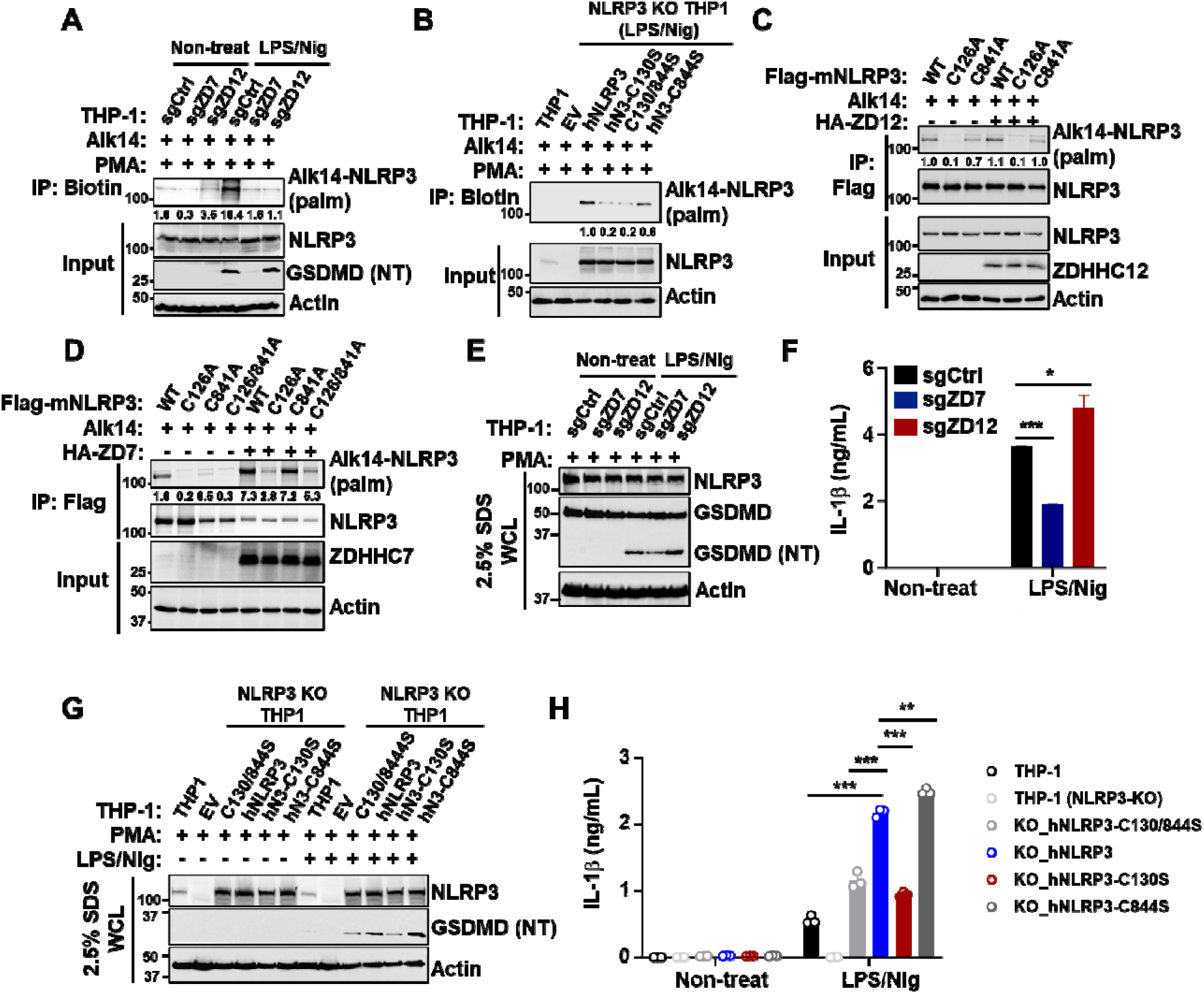
ZDHHC7 promotes NLRP3 activation while ZDHHC12 inhibits NLRP3. **(A)** Palmitoylation of NLRP3 in *ZDHHC7*-deleted (sgZD7), *ZDHHC12*-deleted (sgZD12), or control (sgCtrl) THP-1 cells that were transduced with Cas9 nuclease, primed with PMA, and activated with LPS and 10 μM nigericin for 1 h. NLRP3 palmitoylation level was quantified and normalized by NLRP3 protein level in the input samples. **(B)** Palmitoylation of NLRP3 in wildtype THP-1 or NLRP3-KO THP-1 cells that were reconstituted with NLRP3 WT or cysteine to serine mutants (CS). Cells were primed with PMA, and activated with LPS and nigericin for 1 h as indicated. NLRP3 palmitoylation level was quantified as above. **(C)** Palmitoylation of NLRP3 WT or cysteine to alanine mutants (CA) expressed in HEK 293T cells with ZDHHC12 co-expression. Palmitoylation was detected by Alk14 labeling. NLRP3 palmitoylation level was quantified as above. **(D)** Palmitoylation of NLRP3 WT or cysteine to alanin mutants (CA) in HEK 293T cells with ZDHHC7 co-expression. Palmitoylation was detected by Alk14 labeling. NLRP3 palmitoylation level was quantified as above. **(E)** Immunoblot analysis of NLRP3, GSDMD, and cleaved GSDMD (NT) in the whole cell lysate of *ZDHHC7*-KO (sgZD7), *ZDHHC12*-KO (sgZD12), or control (sgCtrl) THP-1 cells that were transduced with Cas9 nuclease, primed with PMA and LPS, and activated with nigericin (Nig) for 1 h. Cells were lysed with lysis buffer containing 2.5% SDS to get total proteins. **(F)** ELISA of human IL-1β in the cell culture media of THP-1 cells in (**E**). **(G)** Immunoblot analysis of NLRP3 and cleaved GSDMD (NT) in the whole cell lysate of WT or NLRP3-KO THP-1 cells that were reconstituted with NLRP3 WT or cysteine to serine mutant (CS), primed with PMA and LPS, and activated with nigericin for 1h. Cells were lysed with lysis buffer containing 2.5% SDS to get total proteins. **(H)** ELISA of human IL-1β in the cell culture medium in (G). Data with error bars are mean ± SEM. *p < 0.05, **p < 0.01, ***p < 0.001 as determined by the unpaired Student’s t test.

Next, we sought to evaluate the difference between NLRP3 Cys126 and Cys841 (mouse NLRP3 number) palmitoylation. Through reconstituting NLRP3 WT and modification sites mutants into the NLRP3-KO THP-1 cells, we found that all the NLRP3 mutants, C130S, C844S, and C130/844S (human NLRP3 number) had decreased *S*-palmitoylation. However, C844S only decreased about 40% of modification level, while C130S decreased 80% of modification, similar with that of C130/844S mutant (80%) (Figure 8B), which indicate that NLRP3 Cys130 palmitoylation occurred more extensively than the modification on Cys844 in human macrophages. We further determined the NLRP3 S-palmitoylation by expressing mouse NLRP3 WT and mutants in HEK 293T cells and found that NLRP3 C126A had much lower modification level than C841A (Figure 8C, D), and C126/841A double mutant did not further decreased the modification level compared to C126A (Figure 8D), which is consistent with the results of human NLRP3 in THP-1 cells. Additionally, NLRP3 *S*-palmitoylation was strongly promoted by ZDHHC7 co-expression (Figure 8D) but only marginally induced by ZDHHC12 (Figure 8C), further confirming that ZDHHC7-mediated Cys126 palmitoylation occurs more extensively than that of ZDHHC12-mediated Cys841 palmitoylation at resting state.

Lastly, we aim to compare the effects of NLRP3 *S*-palmitoylation between these two sites, catalyzed by two different ZDHHCs, on NLRP3 inflammasome regulation. Our data indicated that *ZDHHC7* deletion inhibited, while *ZDHHC12* KO enhanced, GSDMD cleavage and IL-1β secretion in activated THP-1 cells (Figure 8A, E, F), which is consistent with the above results and previous study^52^. In THP-1 cells that were depleted of endogenous human NLRP3, the C130S or C130/844S mutant-reconstituted cells exhibited decreased GSDMD cleavage and IL-1β secretion compared to wildtype NLRP3-reconstituted cells, while C844S-reconsituted THP-1 showed enhanced GSDMD cleavage and IL-1β secretion (Figure 8G, H), suggesting that NLRP3 Cys126 (Cys130 for human protein) palmitoylation promotes, while Cys841 (Cys844 for human protein) palmitoylation inhibits, NLRP3 inflammasome activation in macrophages. These results suggest that NLRP3 activation is precisely regulated by the *S*-palmitoylation on different sites that are catalyzed by different ZDHHCs to tune the inflammasome process.

## Discussion

In this study, we demonstrated that NLRP3 is palmitoylated on Cys126 by palmitoyl-acyltransferase ZDHHC7. This palmitoylation event is important for NLRP3 localization at th TGN and dTGN, as well as ASC recruitment and oligomerization, and thus inflammasome activation (Figure 9). Cys126 is in the NLRP3 linker region between the Pyrin and NACHT domains^44^ and near the polybasic region that contributes to NLRP3 TGN localization^23,24,51^. Our data showed that NLRP3 S-palmitoylation site mutant disrupts its TGN/dTGN localization (Figure 6), and mutation of the polybasic region also abolishes NLRP3 Cys126 S-palmitoylation (Figure S9A-F). We think the likely explanation for this is that the polybasic region is required for transient NLRP3 interaction with TGN-localized ZDHHC7 (Figure S9G, H), and ZDHHC7-catalyzed palmitoylation then stabilizes NLRP3 Golgi localization. Therefore, it appears that both the polybasic residues and Cys126 S-palmitoylation contribute to localization of NLRP3 at the TGN. PtdIns4P on TGN could recruit NLRP3 through the polybasic region, however, considering that PtdIns4P is found in both the cell plasma membrane and Golgi^53^, it is possible that the localization of ZDHHC7 and its palmitoylation of NLRP3 is the key determinant as to where the NLRP3 inflammasome assembles.

**Figure 9.**
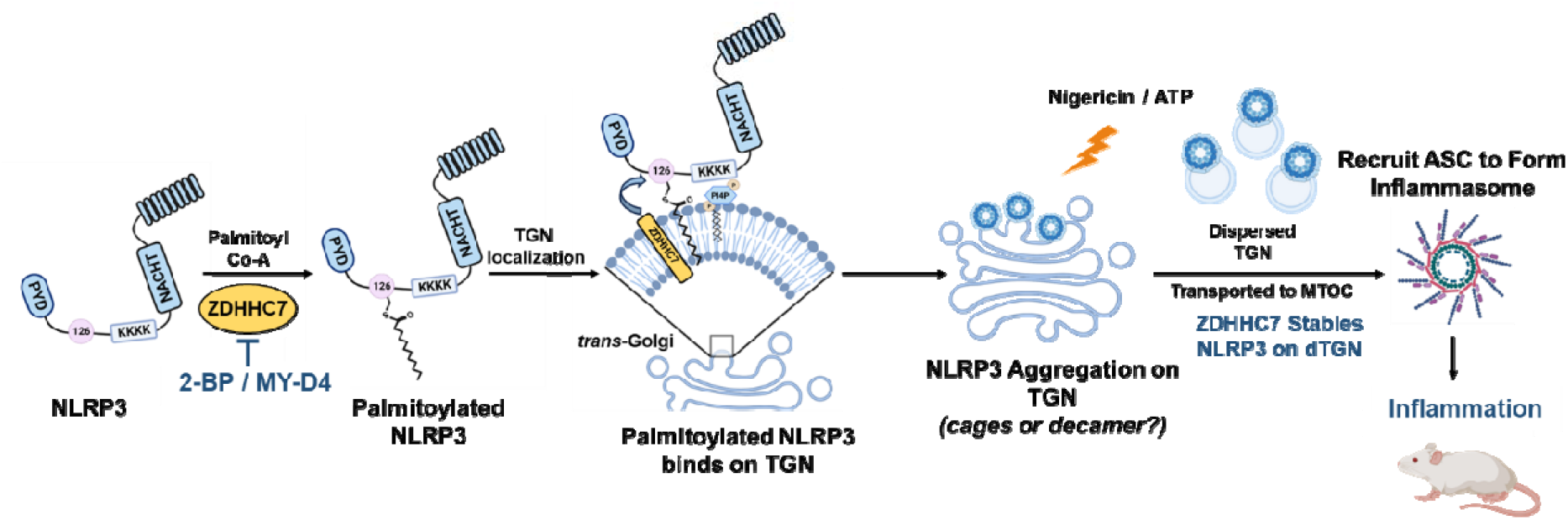
Model for NLRP3 Cys126 palmitoylation regulating NLRP3 inflammasome activation. ZDHHC7-mediated NLRP3 Cys126 palmitoylation promotes NLRP3 *trans*-Golgi network localization and ASC assembly in macrophages, which is essential for NLRP3-mediated inflammasome activation and inflammatory response.

Our data showed that NLRP3 Cys126 palmitoylation is important for ASC recruitment by activated NLRP3 and consequential ASC oligomerization, which is likely a secondary effect of the TGN localization of resting NLRP3 and dTGN localization after activation. Suppressing NLRP3 Cys126 palmitoylation by depleting ZDHHC7, the major palmitoyl-acyltransferase of NLRP3 Cys126 palmitoylation, by pharmacologically inhibiting ZDHHC7 with inhibitors, or by mutating NLRP3 modification site Cys126, dramatically inhibited Caspase-1/GSDMD cleavage and downstream cytokines secretion in macrophages and in mouse models, supporting that ZDHHC7-meidated NLRP3 Cys126 palmitoylation is important for NLRP3 inflammasome activation. Our study shows that NLRP3 is palmitoylated at resting state and the palmitoylation is enhanced in LPS-primed BMDMs or PMA-primed THP-1 cells (Figure 1A, B). Previous work suggested that LPS could significantly accelerate palmitic acid synthesis by inducing the expression of fatty acid synthase (FASN) and the maturation of Sterol Regulatory Element Binding Protein-1c (SREBP-1c) in macrophages^54,55^, which might contribute to the enhanced NLRP3 *S*-palmitoylation in primed macrophages. After activation with ATP or nigericin, NLRP3 *S*-palmitoylation remains (Figure 1A, B, H), suggesting that *S*-palmitoylation may continue to stabilize NLRP3 on dispersed TGN, which facilitates its transport to the MTOC and recruitment of adaptors to mediate inflammasome activation.

Recently, Wang and colleagues reported that NLRP3 is palmitoylated on Cys841 by ZDHHC12 and this negatively regulate NLRP3 inflammasome activation^52^, which is distinct from our study. We compared the ZDHHC7-mediated NLRP3 Cys126 palmitoylation and ZDHHC12-mediatd Cys841 palmitoylation, and found that: (1) NLRP3 Cy126 palmitoylation mediated by ZDHHC7 occurs even at resting state while ZDHHC12-catalyzed NLRP3 Cys841 palmitoylation is induced after NLRP3 activation; (2) NLRP3 Cys126 palmitoylation occurs more extensively than that of Cys841 palmitoylation in macrophages; (3) NLRP3 Cys126 palmitoylation promotes, while Cys841 palmitoylation inhibits, NLRP3 inflammasome activation in macrophages. Therefore, NLRP3 activation is precisely and collaboratively regulated by the ZDHHC7-mediated NLRP3 Cys126 palmitoylation and ZDHHC12-promoted Cys841 palmitoylation in different stages of inflammasome process. Wang and colleagues also found that NLRP3 Cys841 palmitoylation promotes NLRP3 degradation through chaperon-mediated autophagy, however, in our study, NLRP3 total protein level remained the same after 2-BP/MY-D4 incubation, *ZDHHC7*/*ZDHHC12* deletion, or NLRP3 Cys126/Cys841 (human Cys130/Cys844) mutation in macrophages (Figure 2, 8). One possible explanation for the difference is that we used strong detergent (2.5% or 4.0% SDS) to solubilize all proteins, while Wang and co-workers used no detergent or 1% NP-40 lysis buffer, which may not be able to extract membrane-bound NLRP3.

Our study and the previous report point to the following model. In the resting stage, NLRP3 is palmitoylated by ZDHHC7, which promotes NLRP3 TGN localization. After NLRP3 activation, Cys126 palmitoylation persists and ensures NLRP3 localization on dTGN and its transport to MTOC, where NLRP3 recruits ASC to form inflammasome complex. In the late stage of inflammasome activation, ZDHHC12 is induced in macrophages to increase Cys841 palmitoylation and suppress inflammasome. Therefore, NLRP3 S-palmitoylation on different sites have distinct functions, just like different NLRP3 phosphorylation sites mediated by distinct kinase in different stages of inflammasome process could have different regulatory functions^56–60^. Such site-dependent regulatory roles of *S*-palmitoylation will likely be found in other proteins.

ZDHHC7-catalyzed NLRP3 *S*-palmitoylation is conserved in mouse and human macrophages, and the regulatory mechanism contributes to NLRP3 inflammasome activation *in vivo*. Therefore, perturbation of ZDHHC7-mediated NLRP3 Cys126 palmitoylation will lead to suppression of NLRP3 inflammasome and prevent cytokines secretion-mediated inflammation, suggesting that small molecule inhibitors of palmitoyl-acyltransferase ZDHHC7 or that directly targeting NLRP3 Cys126 residue can potentially suppress inflammasome activation and thus may be useful for treating inflammation and autoimmune diseases. The observation that ZDHHC7 and ZDHHC12 have opposing effects on NLRP3 inflammasome activation also suggest that it is beneficial to have ZDHHC7-specific inhibitors for treating inflammasome hyperactivation in human autoimmune diseases.

## Methods

### Mice

The mouse strains used in this study were described previously^26^. Namely, B6.129P2(FVB)-Zdhhc7tm1.2Lusc/Mmmh, RRID: MMRRC_043511-MU, was ordered from the Mutant Mouse Resource and Research Center (MMRRC) at the University of Missouri. *Zdhhc3*, *Zdhhc6*, *Zdhhc9* knock-out mice, and *Nlrp3*-C126A knock-in mice were generated in this study editing with CRISPR/Cas9 technology. All the mice were kept in a specific-pathogen-free facility and all animal experiments were approved by Cornell University’s Institutional Animal Care and Use Committee.

### Plasmids, antibodies, and reagents

pcDNA3-N-Flag-NLRP3 plasmid encoding mouse NLRP3 (Addgene plasmid # 75127) was a gift from Bruce Beutler^61^. pcDNA3-Myc-ASC plasmid encoding human ASC (Addgene plasmid # 73952) were gifts from Christian Stehlik^62^. Plasmids encoding human HA-ASC and NLRP3-GFP were obtained from GenScript. Flag-NLRP3 or NLRP3-GFP Cys to Ser (CS) mutants were constructed by site-directed mutagenesis. Plasmids encoding murine ZDHHC1-23 or the enzymatically inactive mutant ZDHHS7 were kind gifts from Dr. Masaki Fukata, National Institutes of Natural Sciences of Japan.

Immunoblot antibodies for NLRP3 (D4D8T), IL-1β (D4T2D), GSDMD (E7H9G), ASC/TMS1 (D2W8U), phospho-p65 (Ser536, 93H1), and HA-Tag (C29F4) were ordered from Cell Signaling Technology. Antibodies of Caspase-1 (Casper-1) and NLRP3 (Cryo-2) were purchased from AdipoGen. Antibodies for GSDMD (EPR19828) and ZDHHC7 were ordered from Abcam. Antibodies of β-actin (C4) for immunoblot and TGN38 (B-6) for immunofluorescence staining were purchased from Santa Cruz Biotechnology. Antibody for Flag-Tag (A8592) was ordered from Sigma.

2-Bromopalmitic acid (2-BP, CAS: 18263-25-7), lipopolysaccharide (LPS) of *Escherichia coli* O111:B4 or O127:B8, and Phorbol 12-myristate 13-acetate (PMA, CAS: 16561-29-8) were obtained from Sigma. Adenosine 5’-triphosphate disodium salt (ATP, CAS: 987-65-5), poly(dA:dT), and nigericin (Nig, CAS: 28643-80-3) were from InvivoGen. Cross linker disuccinimidyl suberate (DSS) was from ThermoFisher Scientific. Click chemistry reagent TAMRA-azide was from Lumiprobe, tris [(1-benzyl-1H-1,2,3-triazol-4-yl) methyl] amine (TBTA) and CuSO_4_ were from TCI Chemicals, tris(2-carboxyethyl) phosphine HCl (TCEP hydrochloride) was from Millipore. Alkyne 14 (Alk14), 2-BP-Alk and MY-D4 were synthesized in our laboratory. For determination of cell culture media or serum cytokines, mouse IL-1β/IL-1F2 Quantikine ELISA kit, mouse IL-6 Quantikine ELISA kit, mouse IL-18/IL-1F4 ELISA kit, mouse TNF-alpha Quantikine ELISA Kit, mouse HMGB1/HMG-1 ELISA kit, and human IL-1β/IL-1F2 Quantikine ELISA kit were ordered from R&D. Control siRNA-A (sc-37007), siRNA targeting human *ZDHHC7* gene (sc-93249) and mouse *Zdhhc7* gene (sc-155507) were purchased from Santa Cruz Biotechnology.

### Cell culture, transfection, and co-immunoprecipitation

Human HEK293T cells were grown in Dulbecco’s modified Eagle’s medium (DMEM, Gibco) supplemented with 10 % fetal bovine serum (FBS, Gibco), 100 units/mL Penicillin, 100 µg/mL Streptomycin, and 250 ng/mL Amphotericin B. DHHC7 knockout HEK293T was obtained as previously described^26,63^, western blot and quantitative real-time PCR (Q-PCR) were used to confirm the knockout of DHHC7. Human monocytic THP-1 and THP-1 cells with NLRP3 knockout (NLRP3-KO THP-1 cells) were cultured in Roswell Park Memorial Institute 1640 medium (RPMI 1640, Gibco) containing 10 % heat-inactivated FBS, 100 units/mL Penicillin, 100 µg/mL Streptomycin, and 250 ng/mL Amphotericin B.

For HEK293T cells transfection, polyethylenimine hydrochloride (PEI, Polysciences) was used as the transfection reagent. Briefly, HEK293T cells were seeded one day before transfection in 10-cm dishes with DMEM media containing 10% FBS. The next day plasmids encoding indicated proteins were diluted with Opti-MEM™ I Reduced Serum Medium (Thermofisher Scientific) and PEI was added at a ratio of 1:3 for DNA: PEI, then the DNA-PEI mixture was incubated at room temperature for 30 min and added into the HEK293T dishes in a drop-wise manner. After 36-48 h, cells were collected or treated for further analysis.

siRNA transfection in THP-1 cells or NLRP3-KO THP-1 cells was performed with transfection reagent Fugene SI according to manufacturer’s instruction. Briefly, THP-1 cells were seeded in 12-well plate and differentiated with 10 ng/mL PMA. The second day, cell media was changed with RPMI 1640 containing 5% heatinactivated FBS. 10 pmol RNAi duplex was diluted into 50 μL Opti-MEM™ I Reduced Serum Medium, 3 μL Fugene SI was diluted into another 50 μL Opti-MEM™ I Reduced Serum Medium, the diluted RNAi duplex and diluted Fugene SI were combined and incubated at room temperature for 15 min, before RNAi-Fugene mixture was added into the cell wells in a drop-wise manner. Cells were incubated for 36-48 h and treated for further analysis.

For co-immunoprecipitation (Co-IP) of Flag-NLRP3 with ASC expressed in HEK293T cells, after transfection cells were lysed with RIPA buffer (50 mM Tris-HCl, pH 7.4, 150 mM NaCl, 1% NP-40, 0.5% deoxylcholate sodium, 0.1% SDS) containing Protease Inhibitor Cocktail (Sigma, Cat. P8849) and then passed through a 21-G needle 20 times before centrifugation at 9000×g for 5 min. Flag-NLRP3 was immunoprecipitated using anti-FLAG M2 agarose beads (Sigma). Interacting proteins of Flag-NLRP3 were analyzed by immunoblotting analysis of the IP samples, cell lysates were analyzed to detect input proteins and loading controls. For Co-IP of endogenous NLRP3 with ZDHHC7 and ASC, mouse macrophage J774A.1, BMDMs and human macrophage THP-1 cells were lysed with RIPA buffer containing Protease Inhibitor Cocktail and nuclease (Pierce, Cat.88702) after LPS priming or with inflammasome activation, cell lysate was passed through a 21-G needle 20 times before centrifugation at 3000×g for 5 min, supernatant was incubated with anti-NLRP3 or anti-ZDHHC7 antibody as indicated and Protein A/G conjugated agarose beads (Santa Cruz) to immunoprecipitate endogenous NLRP3 or ZDHHC7 proteins, interacting proteins were accessed with immunoblotting analysis of the IP samples and cell lysate were determined to detect input samples and loading controls.

### THP-1 stable cell line construction

To delete *ZDHHC7* or *ZDHHC12* in THP-1 cells, we constructed the THP-1 cells expressing Cas9 nuclease. Vector expressing Cas9 nuclease (GeneCopoeia) was transfected into HEK293T cells with packaging vector psPAX2 and envelope vector pMD2.G, 48 h later the cell culture media containing lentiviral particles was collected and filtered with 0.45 μm filters. The filtered media was then used for lentivirus transduction by infecting THP-1 cells with 8 µg/mL polybrene and incubated for 48 h before changing with fresh media and culturing for another 48 h. Infected THP-1 cells were selected by incubation with 400 ng/mL G-418 for 48 h, remaining live cells were grown up and validated for Cas9 expression. Next, we constructed the THP-1 cells with *ZDHHC7* and *ZDHHC12* KO using THP-1-Cas9 cells. Vectors expressing sgRNA targeting *ZDHHC7* or *ZDHHC12* (GeneCopoeia) were transfected into HEK293T cells along with psPAX2 and pMD2.G, filtered cell culture media containing lentiviral particles was used for transduction by infection THP-1-Cas9 cells, infected cells were selected by incubation with 1 µg/mL puromycin for 48 h, remaining live cells were grown up, validated for ZDHHC7 or ZDHHC12 expression by immunoblotting analysis and Q-PCR, and used for further assays. Sequences of sgRNA targeting *ZDHHC7* include: 5’-TCATGCAGCCATCAGGACAC-3’, 5’-CCGGTCAGCCACGTCAGCCT-3’, 5’-AGTCACCACGAAGTCTGCAT-3’. Sequences of sgRNA targeting *ZDHHC12* include: 5’-CCTGGGGTCCTGGTGCGGAC-3’, 5’-GCTGACCTGGGGAATCACGC-3’, 5’-GTGCTCTTCCTGCACGATAC-3’.

For the NLRP3-reconstituted THP-1 stable cell line construction, WT and mutated NLRP3 cDNA were constructed into pCDH backbone and transfected into HEK293T cells with psPAX2 and pMD2.G, filtered cell culture media containing lentiviral particles was used for transduction by infection NLRP3-KO THP-1 cells, infected cells were selected by incubation with 1 µg/mL puromycin for 48 h, remaining live cells were grown up, validated for NLRP3 expression and relevant sequences, and used for further assays.

### Macrophage preparation and stimulation

Mouse bone-marrow-derived macrophages (BMDMs) were prepared as described previously^64,65^. Briefly, femur and tibia from 8-10 weeks aged mice were used to isolate BM cells, then BM cells were cultured in 10-cm petri dishes with DMEM media containing 20% FBS, 20 ng/mL M-CSF, 100 units/mL Penicillin, 100 µg/mL Streptomycin, and 250 ng/mL Amphotericin B. After culturing for five days, attached cells were detached with EDTA-PBS digestion buffer (PBS containing 10 mM EDTA and 2% FBS) and seeded into culture plates for further assays.

Human peripheral blood mononuclear cells (PBMC)-derived primary macrophages were prepared as described previously^64,66^. Briefly, PBMCs ordered from ATCC (PCS-800-011) were thawed and cultured with RPMI 1640 media containing 10% FBS, 2 mM L-glutamine, 50 ng/mL M-CSF, and 25 ng/mL IL-10 in 12-well plates. After culturing for seven days, floating cells were washed away, the attached cells were mature macrophages and used for further experiments.

For NLRP3 inflammasome activation, mouse BMDMs, J774A.1 cells, and human PBMC-derived macrophages were cultured in DMEM media (for BMDMs and J774A.1 cells) or RPMI 1640 media (for human PBMC-derived macrophages) containing 10% FBS and primed with 200 ng/mL LPS for 4 h before removing LPS. 5 mM ATP or 10 μM nigericin were added to the cells and incubated for 45 min or 1 h, respectively. THP-1 cells or NLRP3-KO THP-1 cells were cultured in RPMI 1640 media containing 10% FBS and differentiated with 10 ng/mL PMA for 24 h before PMA removal. Then 200 ng/mL LPS was incubated for 4 h before LPS removal, 10 μM nigericin was added to the cells and incubated for another 1 h. Macrophage whole cell lysate and cell culture media were collected for immunoblotting analysis and ELISA assay. For 2-BP treatment, 5 μM or 10 μM 2-BP was used after LPS priming and removing, and together with ATP or nigericin incubation (J774A.1 cells, BMDMs and PBMC-derived macrophages), or 1 h before nigericin incubation (THP-1 and NLRP3-KO THP-1 cells).

For non-canonical inflammasome activation, mouse BMDMs were seeded in 12-well plate and primed with 200 ng/mL LPS for 4 h before removing LPS, then cells were transfected with 1 μg LPS using Lipofectamine 3000 Transfection Reagent (Thermofisher Scientific) and incubated for 6 h. For AIM2 inflammasome activation, LPS-primed mouse BMDMs were transfected with 2 μg poly(dA:dT) (InvivoGen) using Lipofectamine 3000 and incubated for 6 h.

Macrophages whole cell lysate was obtained by boiling with 2.5% SDS loading buffer after removing cell culture media and washing with PBS. This procedure was used to make sure that we extract the total proteins in cells, even those that were in aggregates and might be insoluble in other detergents.

### ASC oligomerization assay

ASC oligomerization determination was modified from a previous protocol^67^. After stimulation with ATP or nigericin for 30 min, macrophages were washed with cold PBS and then scraped off the plate and resuspended in 1% NP-40 of HEPES buffer. The cell suspension was passed through a 21-G needle 20 times and centrifuged at 4 ℃, 200 × g for 5 min. The supernatant was transferred into a new tube and centrifuged at 4 ℃, 5000 × g for 10 min. The new supernatant was kept for immunoblotting as loading control and the pellet was washed with PBS twice and cross-linked with 2 mM DSS for 30 min at room temperature. After centrifuging at 4 ℃, 5000 × g for 10 min, the pellet was resuspended with 30 μL SDS-PAGE protein loading buffer and determined by immunoblotting analysis.

### Click chemistry to detect S-palmitoylation

Click chemistry was used for *S*-palmitoylation determination in this study, following the protocol as previously described^26,27^. After plasmids transfection and 50 μM Alk14 incubation for 6 h in HEK293T cells, Flag-NLRP3 was immunoprecipitated with anti-Flag M2 resin, EGFP-NLRP3 was immunoprecipitated with GFP-Trap (Chromotek). The agarose beads were washed with IP washing buffer (25 mM Tris-HCl, pH 8.0, 150 mM NaCl, 0.2% NP-40) three times and resuspended with 20 μL of IP washing buffer. To this bead suspension, click chemistry reagents (2 mM TAMRA-azide, 10 mM TBTA, 40 mM CuSO_4_, 40 mM TCEP, 1 μL each) were added and incubated at room temperature avoiding light. After 30 mins, the reactions were stopped by adding 10 μL SDS loading buffer and boiled at 95 L for 10 mins. For hydroxylamine treatment, the boiled samples were centrifuged, and supernatants were transferred to a new tube. Hydroxylamine was added to the supernatants to be final concentration of 0.5 M and the samples were boiled for another 5 min. Alk14 label signals were assessed by SDS-PAGE and in-gel fluorescence analysis with ChemiDoc Imagers (Bio-Rad).

For Alk14 labeling of NLRP3 in BMDMs and THP-1 cells, after LPS priming, inflammasome activation and Alk14 incubation, 1×10^7^ BMDMs were lysed with 0.6 mL 4 % SDS buffer (4% SDS, 150 mM NaCl, 50 mM TEA, pH7.4) containing protease inhibitors and nuclease (Pierce), to make sure that all the NLRP3 protein was obtained after cell lysis, including the membrane-bond NLRP3. The protein concentration was measured by BCA Protein Assay Kit (Pierce) and normalized. Click chemistry reagents (5mM Biotin-azide, 2 mM TBTA, 50 mM CuSO_4_, 50 mM TCEP, 30 μL for each) were added and incubated at 37 L avoiding light. After 2 h, the reaction was stopped, and protein was precipitated by methanol and chloroform. After centrifugation at 4 L and 4000 × g for 30 mins, the clear upper phase was removed, and methanol was added to wash the central protein phase. Then the protein pellet was collected with another centrifugation at 4 L and 4000 × g for 30 mins. The protein pellet was resuspended with 200 μL of 2 % SDS buffer (2% SDS and 50 μM EDTA in PBS) and diluted with PBS to 2 mL. Streptavidin agarose beads were added to the sample and incubated at room temperature for 1 h. The beads were collected and washed with IP washing buffer three times. The washed beads were suspended in 30 μL SDS loading buffer and boiled at 95 L for 10 mins. The proteins pull-downed by the streptavidin beads, which were labeled with Alk14, were assessed by immunoblotting analysis with anti-NLRP3 antibody. PBS-diluted protein resuspension without streptavidin beads incubation was accessed with immunoblot to detect the input samples and loading controls.

### Real-time quantitative PCR

Total RNA was extracted by E.Z.N.A.® Total RNA Kit I (Omega Bio-tek) according to manufacturer’s instruction. cDNA was synthesized with SuperScript™ VILO™ cDNA Synthesis Kit (Invitrogen). Real-time quantitative PCR (Q-PCR) was performed in triplicate using 2 × Universal SYBR Green Fast qPCR Mix (ABclonal) by QuantStudio™ 7 Flex Real-Time PCR System. Relative expression of genes was calculated using a standard curve method and normalized to the β*-actin (Actb)*. Primers for Q-PCR are listed in Supplementary Table 1.

### Immunofluorescence imaging

After plasmids transfection or stimuli treatment, cells were washed with cold PBS and fixed with 4% paraformaldehyde solution (4% PFA, Santa Cruz) at 4 ℃ for 15 mins, and permeabilized with 0.2% Triton X-100 in PBS at room temperature for 10 mins. The cells were blocked with 3% bovine serum albumin (BSA) in PBST buffer (PBS containing 0.5% Tween-20) at room temperature for 1 h. Then specific primary antibodies were added at a ratio of 1:200 dilution in 3% BSA-PBST buffer and incubated at 4 ℃ overnight. The cells were washed with PBS three times, and fluorescently conjugated secondary antibodies were used at a ratio of 1:1000 dilution in 3% BSA-PBST buffer and incubated at room temperature for 1 h in dark before PBS washing for another three times. NLRP3-Flag was stained with anti-Flag antibody conjugated with Alexa Fluor 488 at a ratio of 1: 500 dilutions in 3% BSA-PBST buffer and incubated at 4 L overnight. DAPI Fluoromount-G Mounting Medium (Southern Biotech) was used for nucleus staining. The stained cells were photographed with LSM710 Confocal Microscope (Zeiss). Quantification of NLRP3 colocalization with TGN markers, TGN38 or 58K Golgi, was analyzed by Image J with the method of Pearson’s correlation coefficient (PCC)^68^, 4-6 fields of vision from several slides were used for analysis in each group.

### Endotoxic shock mouse model

The procedure of LPS-induced endotoxic shock model was modified as previously^67^. Wildtype or *Zdhhc7* knockout mice with B6.129P2(FVB) background were injected intraperitoneally (i.p.) with 35 mg/kg of body weight LPS (lipopolysaccharides from *Escherichia coli* O111:B4, obtained from Sigma, catalog number: L2630, dissolved in sterile PBS). Eight hours later, the mice were sacrificed, and serum was collected (Figure 6a). For 2-BP administration, wildtype mice were administered DMSO (vehicle control, 2 mL/kg), or 25 mg/kg 2-BP by i.p. injection. After 12 h, 35 mg/kg of LPS was injected by i.p. Eight hours later, mice were sacrificed, and the blood was collected via cardiac puncture to prepare serum. IL-1β, IL-6, IL-18, TNFα, and HMGB-1 concentrations in mouse serum were detected by ELISA Kits (R&D) following the manufacturer’s instruction.

### Mono sodium urate crystal (MSU)-induced Peritonitis

The procedure of MSU-induced peritonitis model was modified as previously^67^. Wildtype or *Zdhhc7* knockout mice with B6.129P2(FVB) background were injected intraperitoneally (i.p.) with 1 mg/kg of body weight LPS (Sigma, L2630, in 200 uL sterile PBS) for 3 h and then with 200 μl MSU suspension (i.p., 10 mg/ml, in sterile PBS) for each mouse using a 1-ml syringe, 6 hours after MSU administration the mice were sacrificed, blood was collected via cardiac puncture to prepare serum. Then, inject 3 ml PBS i.p. into each mouse using a 5-ml syringe and collect abdominal contents by peritoneal lavage. Cell numbers of peritoneal exudate cells (PEC) were counted with Bio-Rad TC20 Automated Cell Counter, infiltrated neutrophils and macrophages in peritoneal lavage fluids were determined with flow cytometry by staining and gating with CD45^+^CD11b^+^Gr-1^+^ (neutrophils) or CD45^+^CD11b^+^F4/80^+^ (macrophages) and analyzed with Thermo Fisher Attune NxT Analyzer.

### Statistical analysis

Data are shown as meanL±LSEM, all presented data are representative results of at least three independent repetition, unless specifically indicated. Statistical analysis was performed with GraphPad Prism 9 (Graph-Pad Software), statistics were analyzed using two-tailed Student’s t-test or one-way or two-way ANOVA as indicated. Differences of data were considered to be significant at pL≤L0.05 and are indicated by *, those at pL≤L0.01 are indicated by **, and those at pL≤L0.001 are indicated by ***.

## Acknowledgement

We thank Dr. Bernhard Luscher for providing the DHHC7 knockout strain, B6.129P2(FVB)-Zdhhc7tm1.2Lusc/Mmmh, RRID: MMRRC_043511-MU, which was obtained from the Mutant Mouse Resource and Research Center (MMRRC) at the University of Missouri. We thank Dr. Bruce Beutler for providing pcDNA3-N-Flag-NLRP3 plasmid (Addgene plasmid # 75127), and Dr. Christian Stehlik for providing pcDNA3-Myc-ASC plasmid (Addgene plasmid # 73952). The work is supported by HHMI and Cornell University. Dr. Rob Munroe and Dr. Chris Abratte at the STEM Cell and Transgenic Mouse Facility of Cornell University (supported by NYSTEM) for generating the CRISPR-edited NLRP3 knockout and C126A mutation mice, Dr. Johanna Dela Cruz at the Cornell Institute of Biotechnology’s Imaging Facility for help with immunofluorescence imaging, with NIH 1S10RR025502 funding for the shared Zeiss LSM 710 Confocal Microscope.

## Author Contribution

T.Y. designed and performed all the experiments except those noted below. D.H. designed and performed a variety of experiments to investigate the interaction of NLRP3 with ZDHHC7, compare the function of ZDHHC7 and ZDHHC12, verify the impact of NLRP3 C126 mutation, and compare the regulation of Cys126 and Cys841 palmitoylation, in inflammasome activation. J.Z. designed and performed experiments to identify the site of NLRP3 cysteine palmitoylation and study its function. X.L., D.H. and T.Y. designed and performed the mice study. W.K.G. designed and purified ZDHHC7 proteins. Q.Z. and T.Y. designed and conducted mass spectrometry to identify NLRP3 palmitoylation sites. M.Y. designed and carried out the chemical synthesis of Alk14, MY-D4, 2-BP-Alk. D.C.C. contributed to the genotyping involved in NLRP3 knockout and C126A mouse. M.E.L. provided all the ZDHHC plasmids, directed the *in vitro* DHHC7 assay, and revised the manuscript. H.L. directed the studies and provided overall guidance. T.Y. and H.L. wrote the manuscript with input from all the authors. All authors reviewed and approved the manuscript.

