## Supplementary table and figures for "NLRP3 Cys126 palmitoylation by ZDHHC7 Promotes Inflammasome Activation"

**Supplementary Table 1. Sequences of Q-PCR Primers**

| **Genes** | **Primers** | **Sequences (5’ to 3’)** |
| --- | --- | --- |
| Mouse *Il-1b* | Forward | AAGCCTCGTGCTGTCGGACC |
|  | Reverse | TGAGGCCCAAGGCCACAGGT |
| Mouse *Il-18* | Forward | GACAGCCTGTGTTCGAGGATATG |
|  | Reverse | TGTTCTTACAGGAGAGGGTAGAC |
| Mouse *β-actin* | Forward | CGTGAAAAGATGACCCAGATCA |
|  | Reverse | CACAGCCTGGATGGCTACGT |
| Mouse *Zdhhc1* | Forward | GCGCATGTCATTGAAGACCTGC |
|  | Reverse | CACGCAGTTGTTGAGCCACTTG |
| Mouse *Zdhhc2* | Forward | AGTTCTGAGGCGAGCAGCCAAA |
|  | Reverse | CAGACGGAACAATGATGACAGCG |
| Mouse *Zdhhc3* | Forward | TGGTGGGATTCCACTTCCTGCA |
|  | Reverse | GCCTCAAAGCACAGCAGGATGA |
| Mouse *Zdhhc4* | Forward | CCGGTTTGGGCCGGTTC |
|  | Reverse | CAGATAGCAGCTCCGCTTGG |
| Mouse *Zdhhc5* | Forward | ACACACCTCAGCCTGGCTACTA |
|  | Reverse | ATGGCGGCTGATGTGCTACTGC |
| Mouse *Zdhhc6* | Forward | AGTCTGCCAAGCATACAAGGCG |
|  | Reverse | CAACAGGAGGAAGAGCGTGAAC |
| Mouse *Zdhhc7* | Forward | GCTCTGTCTTCGGTTCATGCTC |
|  | Reverse | CTCAAGGCACAGGAAGACCAAC |
| Mouse *Zdhhc8* | Forward | GTGCCCTATCAGTACAGAGGAC |
|  | Reverse | GGTGCTGTCATCTGCCAGAGTA |
| Mouse *Zdhhc9* | Forward | CTGCTGTGAAGTGCTTTGTGGC |
|  | Reverse | TCTGTGGCAACAGGCTACTGCT |
| Mouse *Zdhhc11* | Forward | CATCCAGCAGAGGAGAAAGAGC |
|  | Reverse | TCGGCGAAAGAGTAGACACTGG |
| Mouse *Zdhhc12* | Forward | GTGCTGACCTGGGGAATCAC |
|  | Reverse | CTGCACATTCACGTAGCCA |
| Mouse *Zdhhc13* | Forward | TGGTTCTAGCCTGGACATCCGA |
|  | Reverse | CCATCGCCAAAGCCGAAACTGT |
| Mouse *Zdhhc14* | Forward | ACAGAAGAGGCTATGTCCAGCC |
|  | Reverse | GCTCTGAATGCACTGGTCTTGG |
| Mouse *Zdhhc15* | Forward | AGAGACCTGAGGTCCAGAAGCA |
|  | Reverse | AGACAGAACAGTGATGGCAGCG |
| Mouse *Zdhhc16* | Forward | TGATGCTGCCTTTGAGCCTGTC |
|  | Reverse | ACCGAGTAGGTTCGGAGGATGA |
| Mouse *Zdhhc17* | Forward | CTTCCTTGCCAACAGCGTTGCT |
|  | Reverse | TGAGGTCCAGACTTCCAGTCTC |
| Mouse *Zdhhc18* | Forward | GGAGACGGAACTACCGCTTCTT |
|  | Reverse | GCTGGTGTCTTTTTCAGAGCGG |
| Mouse *Zdhhc19* | Forward | CGAGCGTGTTTGCTGCCTTCAA |
|  | Reverse | AGGTGAGGATGAAGAGTGGTCC |
| Mouse *Zdhhc20* | Forward | GCAAACCAGAGTGACTACGTCAG |
|  | Reverse | CAGCTCCATTCTCTAGCCACTG |
| Mouse *Zdhhc21* | Forward | CTGAGCTGCTTACTTGCTACGC |
|  | Reverse | TGCCCATGAAGGCAGCTAGTCT |
| Mouse *Zdhhc22* | Forward | GCCTACATCTCCGCTGTCCTTT |
|  | Reverse | ATGGCGAACCAGAGGTAGAGCA |
| Mouse *Zdhhc23* | Forward | GGATATGCGGTATCTGTGTACGG |
|  | Reverse | GGTCAGCGATATTCCGTAAACCG |
| Mouse *Zdhhc24* | Forward | TCTACACAGTGGCTCTCCTGCT |
|  | Reverse | AAAAGCAGCCCAGCACCACACA |
| Mouse *Zdhhc25* | Forward | CGTCACACCTACGGACTATGCT |
|  | Reverse | AGTGGCAAGCTGTCCTCGGTAT |
| Human Z*DHHC7* | Forward | CTGACCGGGTCTGGTTCATC |
|  | Reverse | CATGACGAAAGTCACCACGAA |
| Human *β-ACTIN* | Forward | GCAAGCAGGACTATGACGAG |
|  | Reverse | CAAATAAAGCCATGCCAATC |

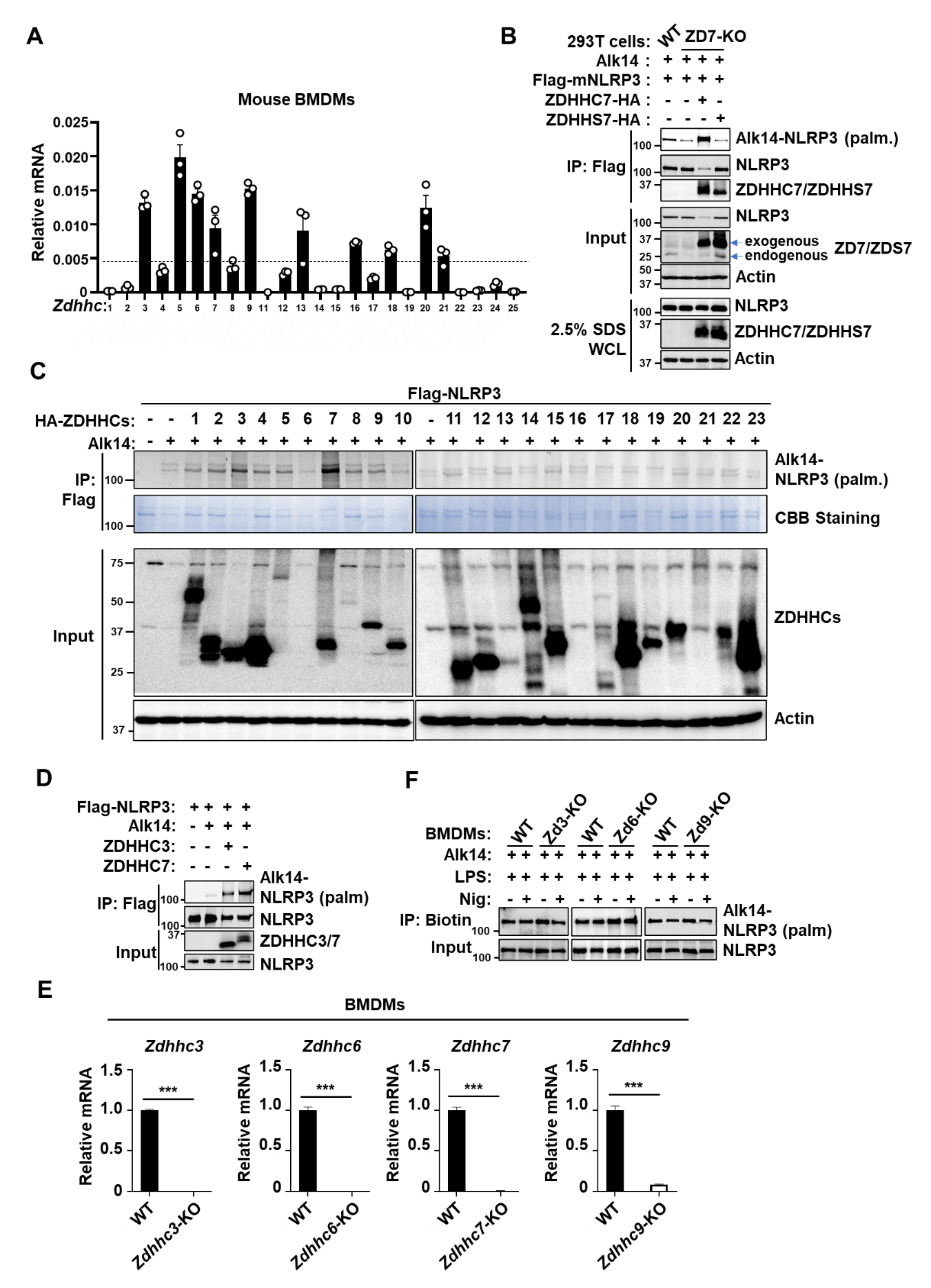

**Figure S1. NLRP3 is palmitoylated by ZDHHC7 in macrophages, related to Figure 1.**

**(A)** Relative mRNA of *Zdhhc* genes in BMDMs determined by quantitative real-time PCR (Q-PCR), mRNA level was normalized to *β-actin*. **(B)** Palmitoylation of Flag-NLRP3 expressed in wildtype (WT) HEK 293T cells, or *ZDHHC7*-deleted (ZD7-KO) HEK 293T cells reconstituted with wildtype ZDHHC7 or its enzymatic DHHC motif mutant (ZDHHS7). Palmitoylation of NLRP3 was detected using Alk14 labeling and in-gel fluorescence. **(C)** NLRP3 palmitoylation determination in HEK293T cells that were transfected with Flag-NLRP3 and each of the palmitoyl-transferases ZDHHCs (ZDHHC1-23) and incubated with 50 μM Alk14 probe, assessed by in-gel fluorescence and immunoblotting analysis. NLRP3 was tagged with Flag (Flag-NLRP3) and ZDHHCs were tagged with HA (HA-DHHCs). **(D)** NLRP3 palmitoylation determination in HEK293T cells that were transfected with Flag-NLRP3, ZDHHC3-HA, or ZDHHC7-HA as indicated. **(E)** Relative mRNA of *Zdhhc3, Zdhhc6, Zdhhc7*, and *Zdhhc9* in BMDMs determined by Q-PCR, showing the relevant *Zdhhc* was knocked-out successfully in BMDMs. **(F)** Palmitoylation of NLRP3 in wildtype (WT) and *Zdhhc3*, *Zdhhc6*, *Zdhhc9*-deleted (Zd3-KO, Zd6-KO, Zd9-KO, respectively) BMDMs by Alk14 labeling and click chemistry assay. BMDMs were incubated with Alk14, LPS and nigericin (Nig) as indicated, cells were then lysed and proteins were conjugated with biotin-azide, labeled proteins were pulled down with streptavidin and blotted for NLRP3. Data with error bars represents mean ± SEM. ∗p < 0.05, ∗∗p < 0.01, ∗∗∗p < 0.001 as determined by unpaired Student’s t test.

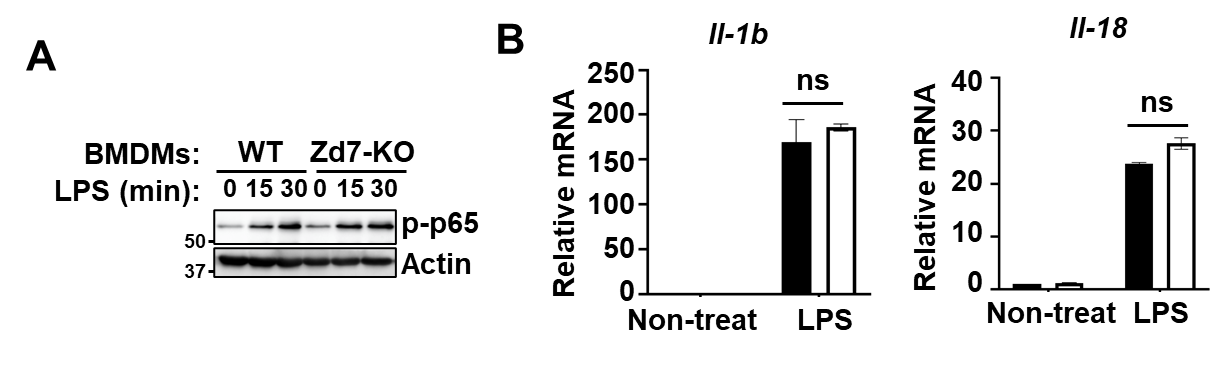

**Figure S2. Zdhhc7 did not regulate NF-κB activation.**

**(A)** Phosphorylation of NF-κB (p65) determined by immunoblotting analysis with anti-phospho-p65 (Ser536) in wildtype (WT) and *Zdhhc7*-deleted (Zd7-KO) BMDMs that were treated with 100 ng/mL LPS for the indicated time. **(B)** Relative mRNA of *Il-1b* and *Il-18* determined by Q-PCR in wildtype (WT) and *Zdhhc7*-deleted (Zd7-KO) BMDMs that were treated with 100 ng/mL LPS for 6 h, mRNA was normalized to *β-actin*. Data with error bars represent mean ± SEM. ∗p < 0.05, ∗∗p < 0.01, ∗∗∗p < 0.001 as determined by unpaired Student’s t test.

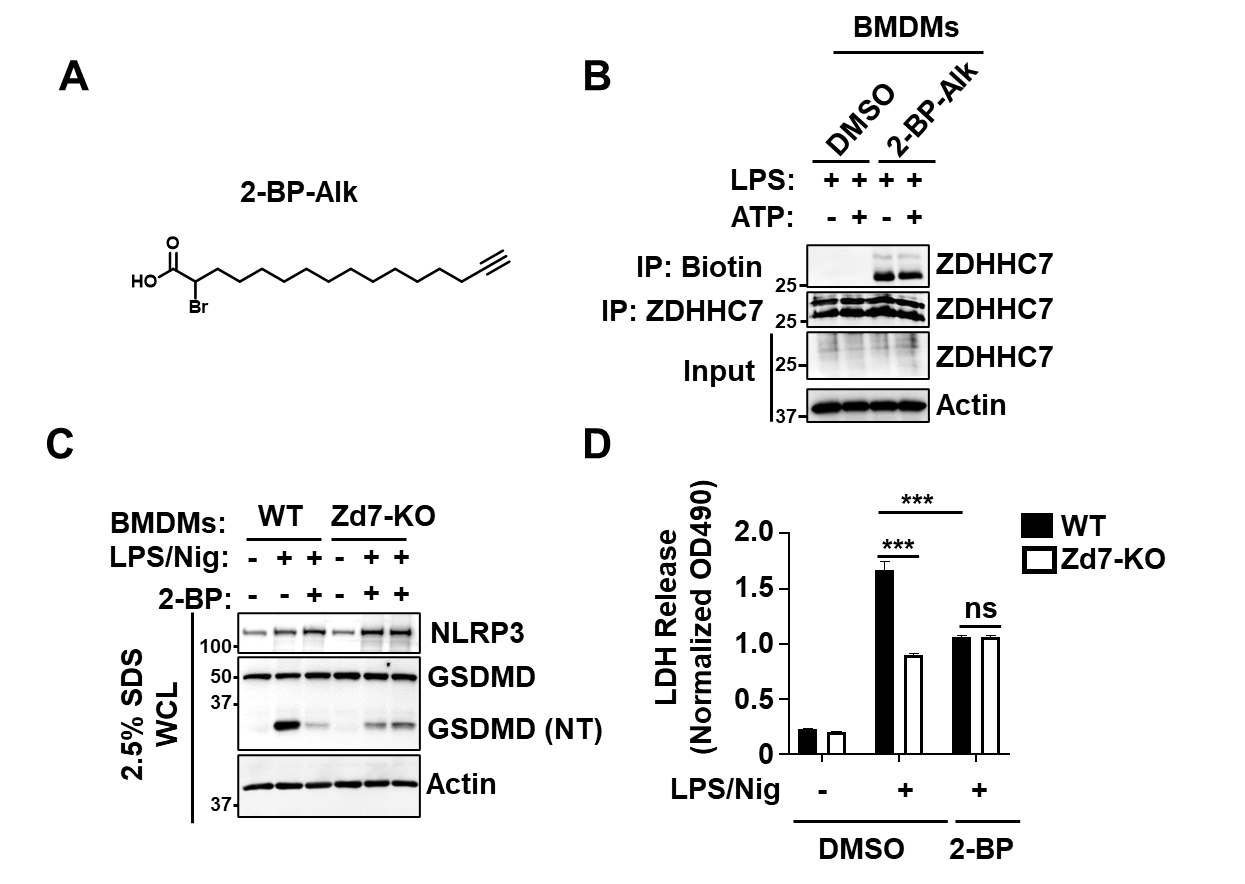

**Figure S3. 2-BP targets ZDHHC7 in BMDMs.**

**(A)** Chemical structure of 2-BP-Alk probe. **(B)** Immunoblot analysis showing 2-BP-Alk covalently bound to ZDHHC7 in LPS-primed and ATP activated BMDMs. 2-BP-Alk was incubated with BMDMs for 6 h, during which LPS was added for 4 h and ATP was added for 30 mins before cell collection. Whole cell lysate was conjugated with biotin-azide and then immunoprecipitated with streptavidin resin, or with anti-ZDHHC7 as control, ZDHHC7 was assessed by immunoblot. **(C)** Immunoblot analysis of NLRP3, GSDMD, cleaved GSDMD (NT) and Actin in wildtype (WT) and *Zdhhc7*-deleted (Zd7-KO) BMDMs that were primed with LPS and activated with nigericin (Nig), with DMSO or 10 μM 2-BP treatment as indicated. **(D)** LDH release assay of BMDMs in cell culture medium showing cell pyroptotic level in (**C**). Data with error bars represent mean ± SEM. ∗p < 0.05, ∗∗p < 0.01, ∗∗∗p < 0.001 as determined by unpaired Student’s t test.

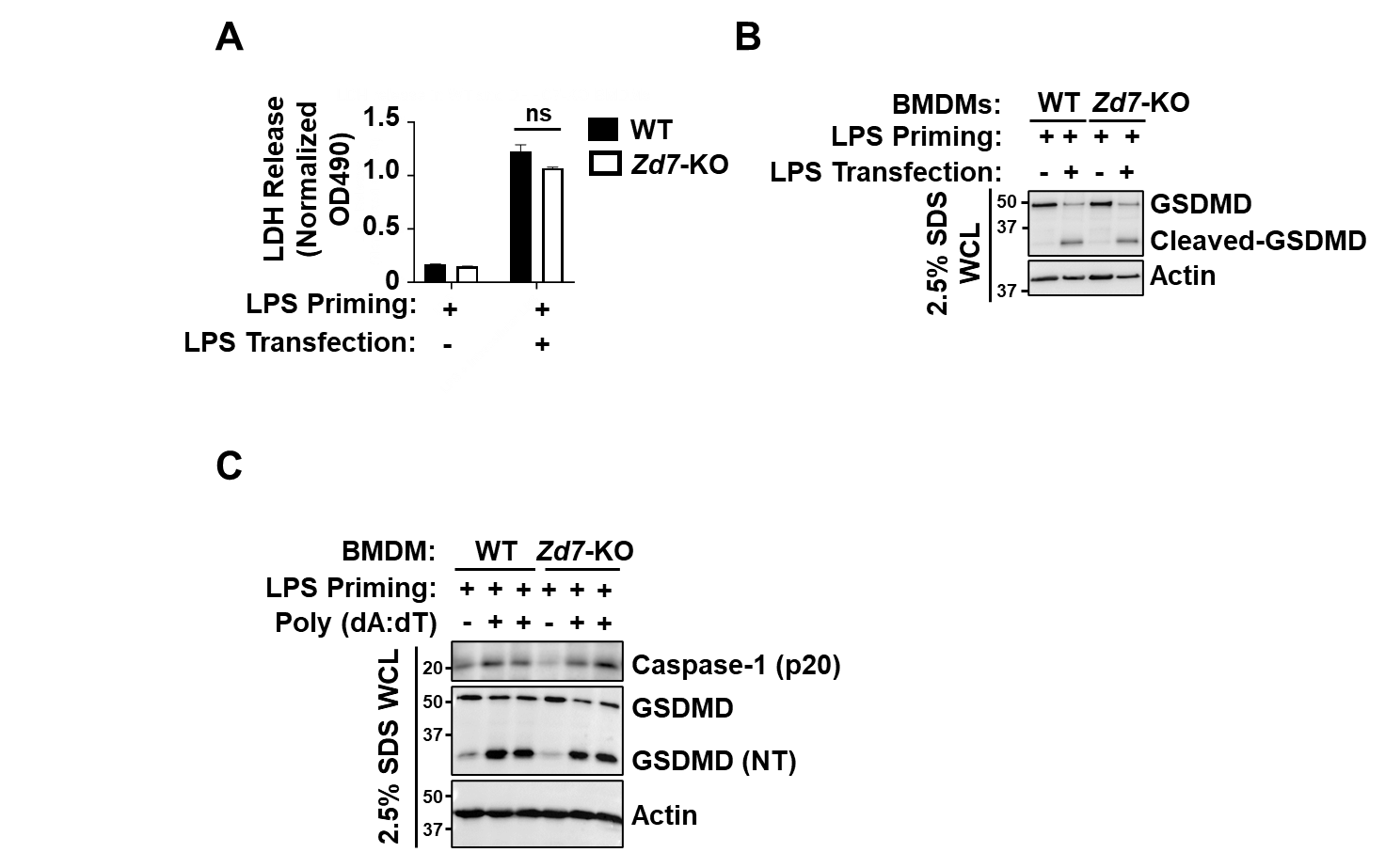

**Figure S4. *Zdhhc7* deletion in macrophage does not regulate non-canonical or AIM2 inflammasome activation.**

**(A)** LDH release assay (pyroptosis) of non-canonical inflammasome activation in wildtype and *Zdhhc7*-KO BMDMs that were primed with 200 ng/mL LPS and activated by LPS transient transfection (1 μg/mL, 6 hours). **(b)** Immunoblotting analysis of GSDMD and GSDMD cleavage (NT) in (**A**). **(C)** Immunoblotting analysis of GSDMD, cleaved Caspase-1 (p20) and cleaved GSDMD (NT) for AIM2 inflammasome activation in wildtype and *Zdhhc7*-KO BMDMs that were primed with 200 ng/mL LPS and activated by poly(dA:dT) transfection (2 μg/mL, 6 hours). Data with error bars represent mean ± SEM. ∗p < 0.05, ∗∗p < 0.01, ∗∗∗p < 0.001 as determined by unpaired Student’s t test.

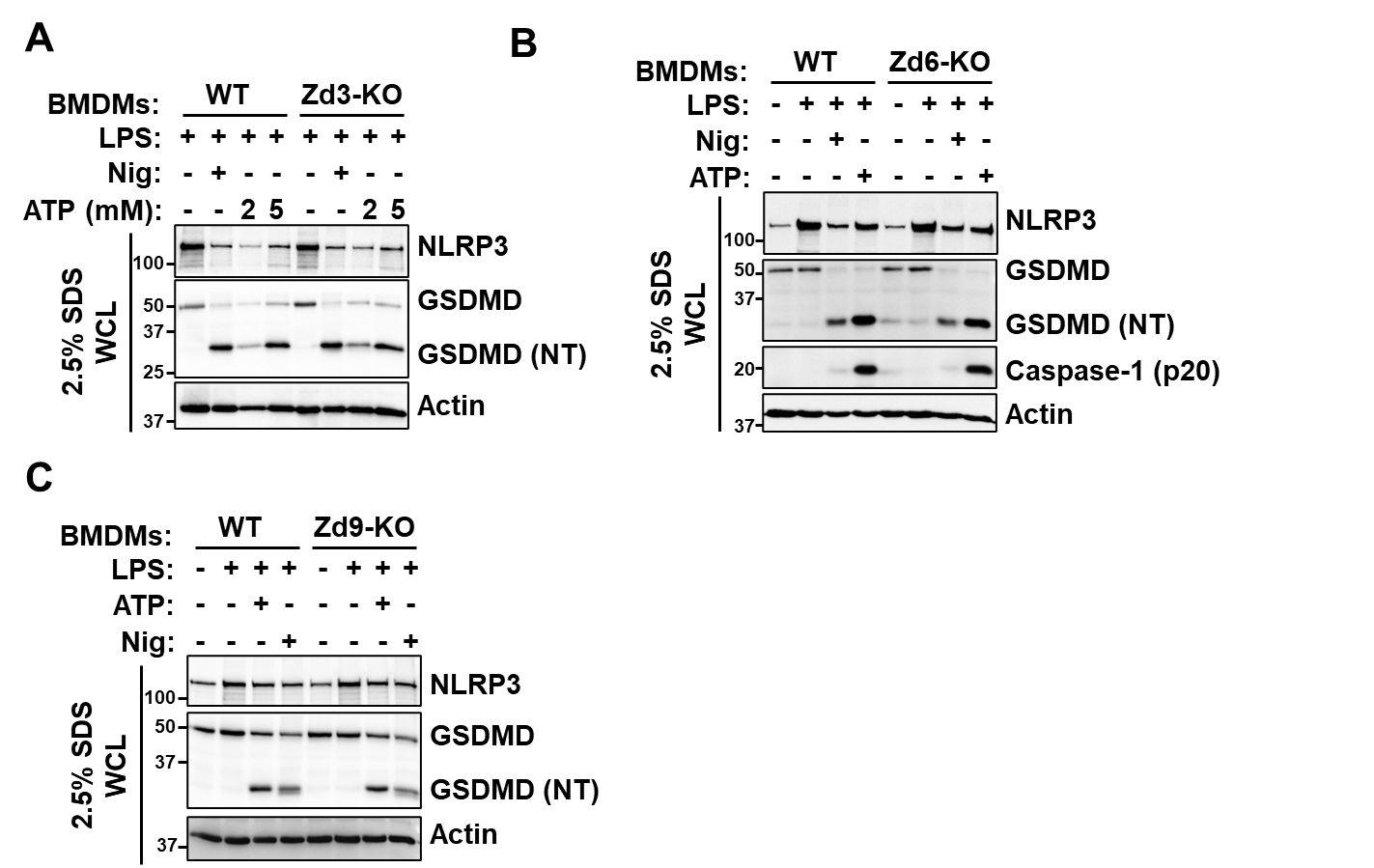

**Figure S5. Deletion of *Zdhhc3*, *Zdhhc6*, or *Zdhhc9* does not affect NLRP3 activation.**

**(A-C)** Immunoblotting analysis of NLRP3, GSDMD and GSDMD cleavage (NT) in whole cell lysate of wildtype (WT), Zdhhc*3*-deleted (Zd3-KO, **A**), *Zdhhc6*-deleted (Zd6-KO, **B**), or *Zdhhc9*-deleted (Zd9-KO, **C**) BMDMs that were primed by LPS and activated with nigericin (Nig) or ATP as indicated. Cells and culture medium were lysed with lysis buffer containing 2.5% SDS to get total proteins.

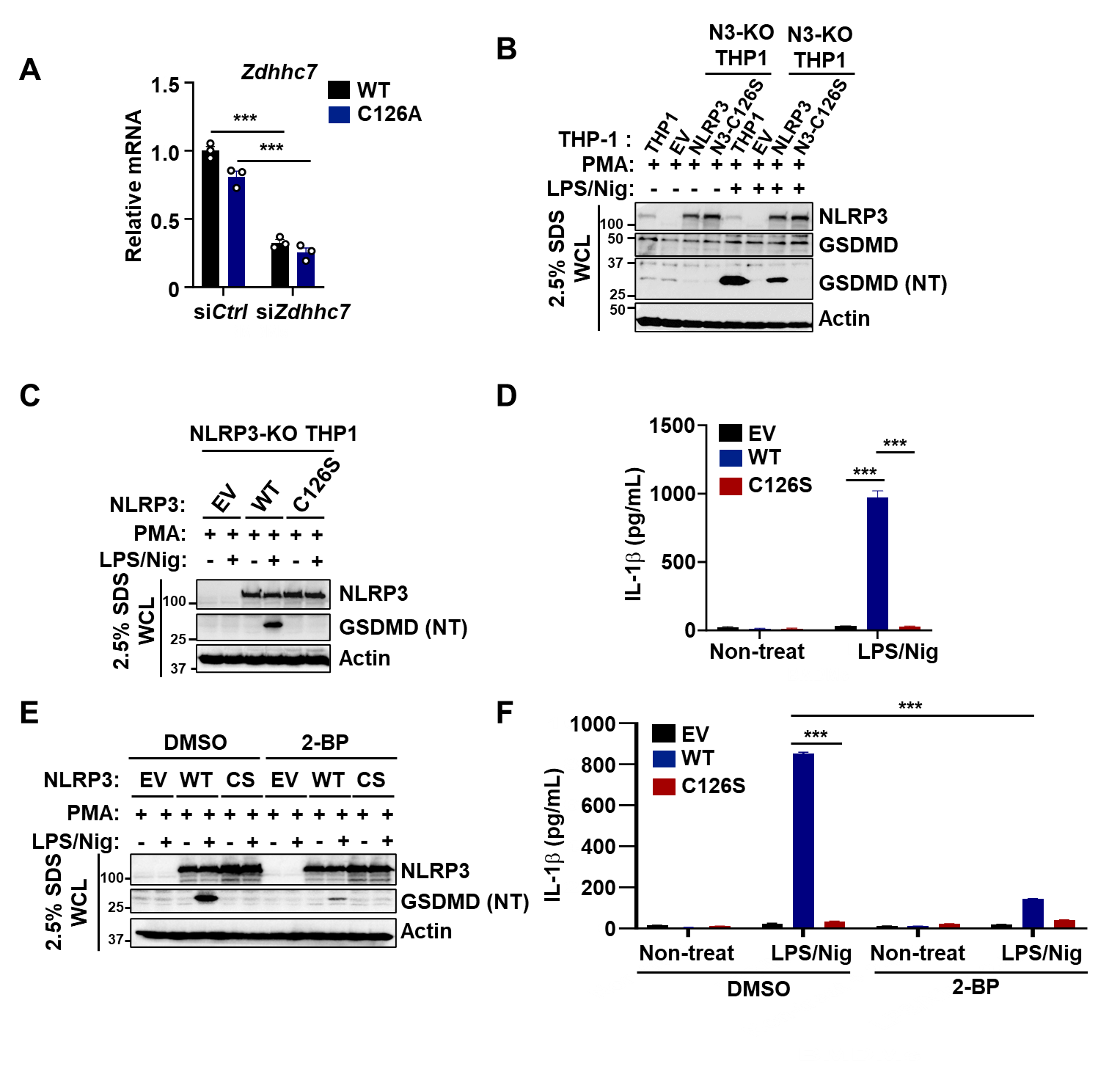

**Figure S6. NLRP3 Cys126 mutant inhibited NLRP3 inflammasome activation in macrophages.**

**(A)** Relative mRNA of *Zdhhc7* determined by Q-PCR in wildtype (WT) and *Nlrp3*-C126A (C126A) BMDMs that were knocked down with either control or *Zdhhc7* siRNA (si*Zd7*) as indicated. mRNA level was normalized to *β-actin*. **(B)** Immunoblot analysis of NLRP3, GSDMD, and cleaved GSDMD (NT) in PMA-primed wildtype or the NLRP3-KO (N3-KO) THP-1 cells that were reconstituted with wildtype or C126S mutant NLRP3 and stimulated with LPS and nigericin as indicated. **(C)** Replication of immunoblot analysis in (**B**). **(D)** ELISA determination of human IL-1β in the cell culture media in (**C**). **(E)** Immunoblot analysis of NLRP3 and GSDMD cleavage (NT) in PMA-primed NLRP3-KO THP-1 cells that were reconstituted with wildtype or C126S mutant NLRP3. Cells were PMA-differentiated, LPS primed, treated with DMSO or 10 μM 2-BP, and stimulated with 10 μM nigericin for 1 hour. **(F)** ELISA determination of human IL-1β in (**E**). Data with error bars represent mean ± SEM. ∗p < 0.05, ∗∗p < 0.01, ∗∗∗p < 0.001 as determined by unpaired Student’s t test.

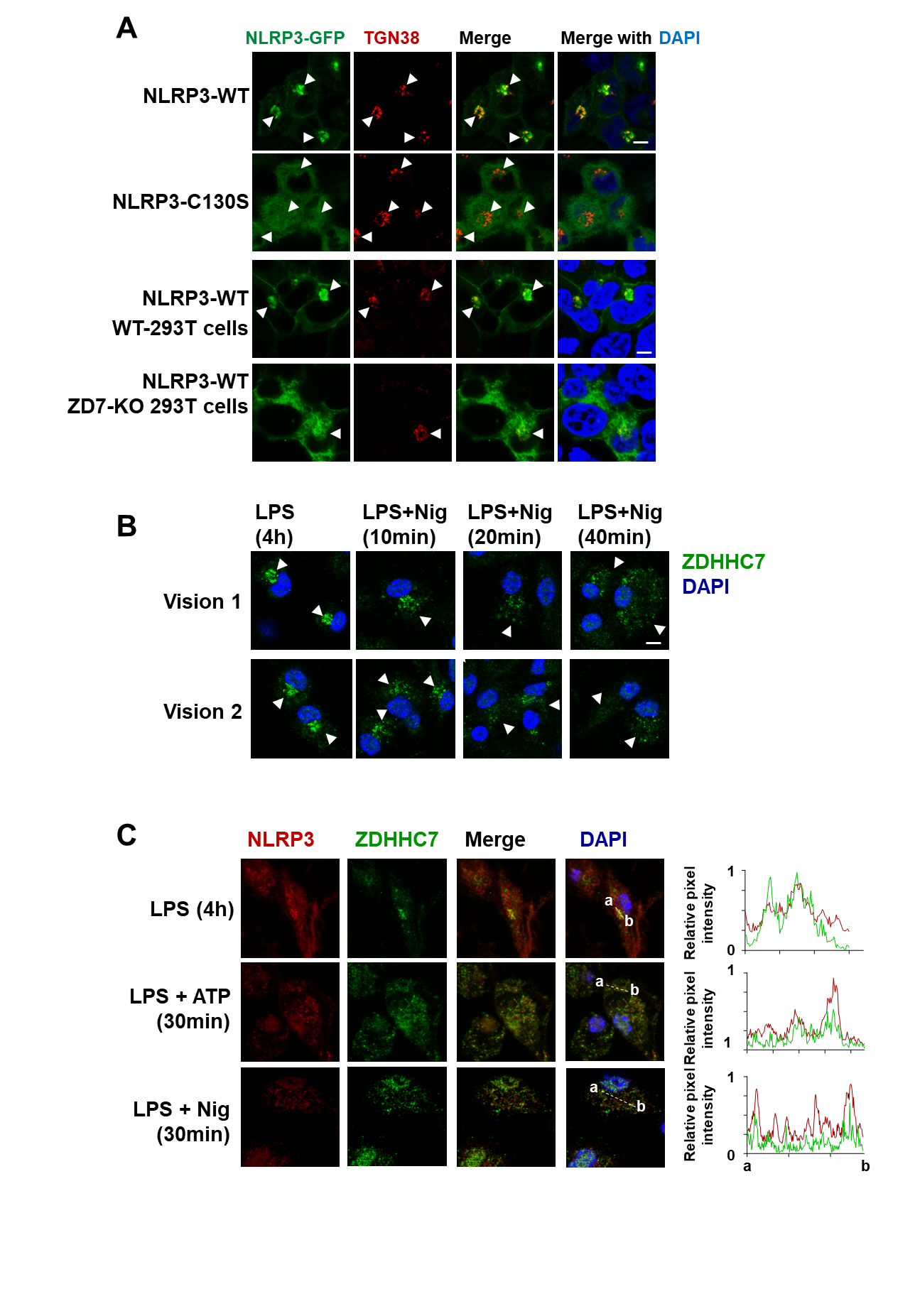

**Figure S7. Human NLRP3 Cys130 S-palmitoylation promotes NLRP3 locating on *trans*-Golgi network.**

**(A)** Representative confocal microscopic images of human wildtype and C130S mutant of human NLRP3 (NLRP3-GFP) in HEK 293T cells, or in *ZDHHC7*-KO (ZD7-KO) HEK 293T cells. WT NLRP3, but not the C130S mutant, colocalized with TGN marker TGN38. WT NLRP3 did not colocalize with TGN marker in *ZDHHC7*-KO cells. NLRP3 localization was shown as green, TGN38 antibody was used to stain *trans*-Golgi network, DAPI was used to stain the nucleus. Scale bar: 5 μm. **(B)** Representative confocal microscopic images of endogenous ZDHHC7 in BMDMs that were primed with 200 ng/ml LPS for 4 hours and activated with 10 μM nigericin as indicated. ZDHHC7 antibody was used to stain ZDHHC7 (green), DAPI was used to stain the nucleus. Scale bar: 5 μm. The images suggested Golgi-localized ZDHHC7 was dispersed during inflammasome activation in BMDMs. **(C)** Representative images showing the locations of endogenous NLRP3 and ZDHHC7 along with the DAPI signal in BMDMs that were primed with 200 ng/ml LPS for 4 hours and activated with 5 mM ATP or 10 μM nigericin (Nig) for 30 min as indicated. Representative curves (right) describe the distribution of relative fluorescence intensities for NLRP3 (red) and ZDHHC7 (green). Scale bar: 5 μm. Data represent at least two independent experiments.

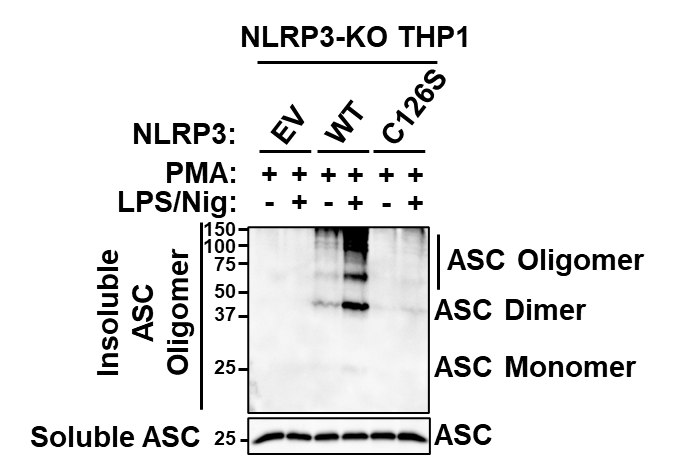

**Figure S8. Cys126 S-palmitoylation promotes ASC oligomerization after inflammasome activation.**

Immunoblot analysis of ASC oligomerization from insoluble protein fraction by DSS cross-linking in PMA-differentiated and LPS-primed THP-1 cells that were reconstituted with wildtype or C126S mutant of NLRP3 and activated with 10 μM nigericin for 1 h.

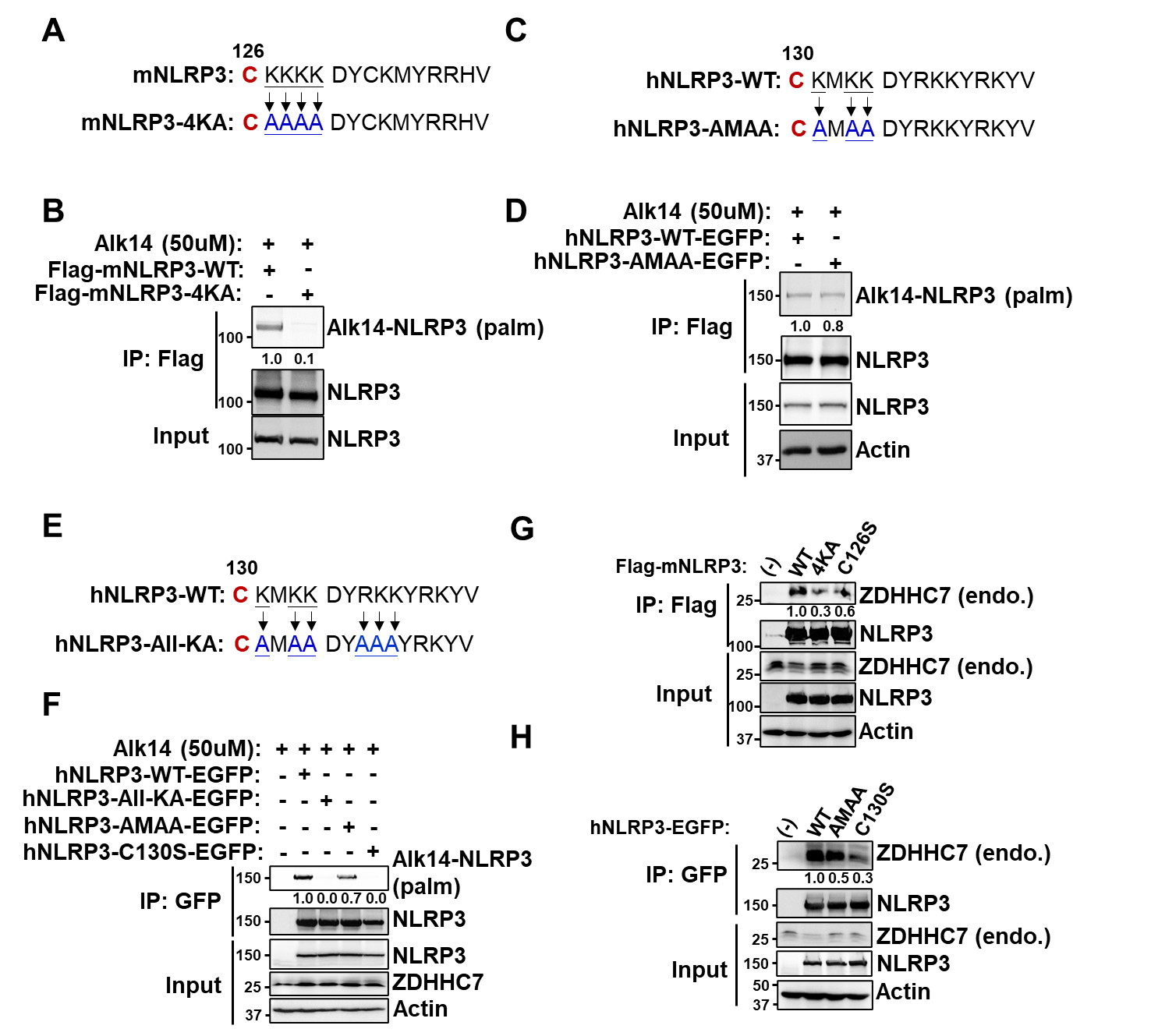

**Figure S9. NLRP3 polybasic region is important for ZDHHC7-catalyzed NLRP3 Cy126 palmitoylation.**

**(A)** Schematic diagram of amino acids sequence for mouse NLRP3 protein polybasic region with lysine mutated to alanine (mNLRP3-4KA), which disrupts NLRP3 TGN localization. **(B)** Palmitoylation of wildtype (WT) or polybasic region demolition mutants (4KA) NLRP3 expressing in HEK 293T cells by Alk14 labeling and click chemistry assay. NLRP3 palmitoylation level was quantified and normalized to NLRP3 protein level of input samples. **(C)** Schematic diagram of amino acids sequence for human NLRP3 protein polybasic region with partially lysine mutated to alanine (hNLRP3-AMAA), which is not sufficient to prevent NLRP3 TGN localization. **(D)** Palmitoylation of human WT or AMAA mutant of NLRP3 in HEK 293T cells detected by Alk14 labeling and click chemistry assay. NLRP3 palmitoylation level was quantified as above. **(E)** Schematic diagram of amino acids sequence for human NLRP3 protein polybasic region with all nearby K/R mutated to alanine (hNLRP3-All-KA), which disrupts NLRP3 TGN localization. **(F)** Palmitoylation of human WT, All-KA, AMAA, or C130S NLRP3 in HEK 293T cells detected by Alk14 labeling and click chemistry assay. NLRP3 palmitoylation level was quantified as above. **(G-H)** Immunoblotting assay of endogenous (endo.) ZDHHC7 interaction with mouse WT or 4KA NLRP3 (**G**), or human WT or AMAA NLRP3 (**H**) in HEK 293T cells. Cell lysate was immunoprecipitated with anti-Flag resin to pull-down NLRP3, endogenous ZDHHC7 was detected by immunoblot with anti-ZDHHC7 in the immunoprecipitation (IP) samples. The Co-IP ratio of ZDHHC7/NLRP3 was calculated and normalized to protein levels in input samples.
